# Attenuated humoral responses in HIV infection after SARS-CoV-2 vaccination are linked to global B cell defects and cellular immune profiles

**DOI:** 10.1101/2022.11.11.516111

**Authors:** Emma Touizer, Aljawharah Alrubbayi, Rosemarie Ford, Noshin Hussain, Pehuén Pereyra Gerber, Hiu-Long Shum, Chloe Rees-Spear, Luke Muir, Ester Gea-Mallorquí, Jakub Kopycinski, Dylan Jankovic, Christopher Pinder, Thomas A Fox, Ian Williams, Claire Mullender, Irfaan Maan, Laura Waters, Margaret Johnson, Sara Madge, Michael Youle, Tristan Barber, Fiona Burns, Sabine Kinloch, Sarah Rowland-Jones, Richard Gilson, Nicholas J Matheson, Emma Morris, Dimitra Peppa, Laura E McCoy

## Abstract

People living with HIV (PLWH) on suppressive antiretroviral therapy (ART) can have residual immune dysfunction and often display poorer responses to vaccination. We assessed in a cohort of PLWH (n=110) and HIV negative controls (n=64) the humoral and spike-specific B-cell responses following 1, 2 or 3 SARS-CoV-2 vaccine doses. PLWH had significantly lower neutralizing antibody (nAb) titers than HIV-negative controls at all studied timepoints. Moreover, their neutralization breadth was reduced with fewer individuals developing a neutralizing response against the Omicron variant (BA.1) relative to controls. We also observed a delayed development of neutralization in PLWH that was underpinned by a reduced frequency of spike-specific memory B cells (MBCs) and pronounced B cell dysfunction. Improved neutralization breadth was seen after the third vaccine dose in PLWH but lower nAb responses persisted and were associated with global, but not spike-specific, MBC dysfunction. In contrast to the inferior antibody responses, SARS-CoV-2 vaccination induced robust T cell responses that cross-recognized variants in PLWH. Strikingly, a subset of PLWH with low or absent neutralization had detectable functional T cell responses. These individuals had reduced numbers of circulating T follicular helper cells and an enriched population of CXCR3^+^CD127^+^CD8^+^ T cells after two doses of SARS-CoV-2 vaccination, which may compensate for sub-optimal serological responses in the event of infection. Therefore, normalisation of B cell homeostasis could improve serological responses to vaccines in PLWH and evaluating T cell immunity could provide a more comprehensive immune status profile in these individuals and others with B cell imbalances.

## INTRODUCTION

People living with HIV (PLWH) appear to be at a higher risk of hospitalisation and worse clinical outcomes from COVID-19 disease, especially in the context of cellular immunosuppression and unsuppressed HIV viral load (Cooper et al., 2020). Although antiretroviral therapy (ART) has dramatically improved life expectancy in PLWH, the persistence of immune dysfunction raises concerns about the overall effectiveness and durability of vaccine responses in this potentially more vulnerable patient group, in line with other immunocompromised groups (Herzog Tzarfati et al., 2201; Kamar et al., 2021). As a result, PLWH were included in priority group 4 and 6 in the UK for earlier COVID-19 vaccination than the general population. The Joint Committee on Vaccination and Immunization (JCVI) advised to invite this patient group for a 4^th^ booster dose (Baskaran et al., 2021; Bertagnolio et al., 2022; Dandachi et al., 2021; Hoffmann et al., 2021; Jcvi, 2022; Noe et al., 2021; Western Cape Department of Health in collaboration with the National Institute for Communicable Diseases, 2021; Yang et al., 2021). Previously, defects have been observed in serological vaccine responses in PLWH. For example after a full course of hepatitis B (Cruciani et al., 2009) or influenza vaccination (George et al., 2015) and long-term responses to vaccination can be shorter-lived in PLWH compared to the general population (Kernéis et al., 2014). We and others have previously shown a failure to mount a robust antibody response following COVID-19 vaccination in advanced HIV infection with low CD4 T cell counts below 200 cells/µl (Hassold et al., 2022; Nault et al., 2021; Noe et al., 2021; Spinelli et al., 2021; Touizer et al., 2021).

Data on vaccine efficacy and immunogenicity in PLWH remains limited (reviewed in (Mullender et al., 2022)), and while there are some conflicting results, meta-analyses (Tamuzi et al., 2022) and recent studies (Woldemeskel et al., 2022) have shown reduced levels of seroconversion and neutralization after a second dose of viral vector vaccine dose in PLWH, with lower CD4 T cell count/viraemia and older age resulting in a more impaired response and more rapid breakthrough infection (Sun et al., 2022). Data after three vaccine doses are scarce, especially of evaluating efficacy against Omicron. However, the data available thus far suggest that the third vaccine dose provides a strong boost to antibody responses regardless of the CD4 T cell count, including in those who had previously not seroconverted (Vergori et al., 2022). Moreover, most studies on SARS-CoV-2 vaccine responses in PLWH to date have mostly focussed on evaluating humoral responses and generated limited data on functional T cell responses (Ogbe et al., 2022b) or cellular profiles of T or B cells. Therefore, it remains unclear what role HIV-associated immune dysfunction plays in serological and cellular outcome after SARS-CoV-2 vaccination.

Inferior serological responses to vaccination in PLWH are most commonly linked to HIV-induced immune destruction of CD4 T cells and imbalance of the CD4:CD8 T cell populations (Fuster et al., 2016; Pallikkuth et al., 2018). Despite effective ART, chronic immune activation in HIV can lead to exhaustion of the adaptive immune system (Fenwick et al., 2019). This can translate into impaired T cell responses, likely limiting T follicular helper (T_FH_) cell help to B cells, resulting in lower serological outputs. There is also substantial evidence for dysfunction/exhaustion in the B cell compartment during chronic infections that may limit antibody responses against the infecting pathogen (Burton et al., 2018; Portugal et al., 2015). This B cell dysfunction persists to a variable degree after HIV viral suppression (Moir and Fauci, 2013, 2017), but how these B cell defects impact serological responses to vaccination has not yet been fully elucidated. Furthermore, there is substantial age-related decline in immune function leading to senescence in both the T and B cell compartments, which may be accelerated in PLWH and could further influence vaccine responses (Nasi et al., 2017).

In this study we have evaluated in a well-curated cohort of PLWH and HIV-negative controls following three SARS-CoV-2 vaccine doses, the relationship between humoral and functional T cell responses against Omicron and other variants of concern (VOC). To achieve this goal, we have assessed how spike-specific memory B cell (MBC) responses, global MBC profiles, CD4 and CD8 T cell phenotypes are linked with serological outcomes in PLWH to better understand which factors may modulate immune responses to vaccination.

## RESULTS

### Lower levels of seroconversion and neutralizing antibodies after SARS-CoV-2 immunization in PLWH without a history of prior COVID-19 disease

Participants were recruited between January 2021 and April 2022 (n=110 PLWH and n=64 HIV-negative controls) as described in **Table 1**. Participants were sampled after 1, 2 or 3 doses of a SARS-CoV-2 vaccine and compared cross-sectionally. In addition, in 53 PLWH and 44 controls, responses were assessed longitudinally where sequential samples were available. SARS-CoV-2 spike-specific IgG were tested for binding against the S1 subunit of the SARS-CoV-2 spike protein in a semi-quantitative ELISA (Ng et al., 2020; Rees-Spear et al., 2021) to determine seropositivity. Neutralizing antibodies (nAbs) were measured against the ancestral vaccine-matched Wuhan Hu-1 SARS-CoV-2 (WT) strain by pseudovirus neutralization (Rees-Spear et al., 2021). Approximately 90% of HIV-negative controls and 80% of PLWH with no prior history of SARS-CoV-2 infection seroconverted. However, while over 82% of controls produced a neutralizing response after one vaccine dose, only 29% of PLWH did so (**Figure 1A**). As described (Reynolds et al., 2021), prior history of SARS-CoV-2 infection was associated with a higher level of seroconversion and the development of nAbs in all individuals at every studied timepoint regardless of HIV status **(Figure 1A)**.

**Table 1:** Cohort Demographics. Cohort demographics, clinical characteristics and number of participants per timepoints for each group. AZ= AZD1222; Moderna= mRNA-1273 Pfizer= BNT162b2.

|  | HIV- (n=64) |  | HIV+ (n=110) |  |
| --- | --- | --- | --- | --- |
|  | SARS-CoV-2-<br>(n=27) | SARS-CoV-2+<br>(n=37) | SARS-CoV-2-<br>(n=65) | SARS-CoV-2+<br>(n=45) |
| **Clinical parameters** |  |  |  |  |
| % female | 76% | 40% | 8% | 17% |
| % BAME | 28% | 24% | 21% | 44% |
| Median age (range) | 33(21-65) | 41(23-66) | 53 (22-93) | 49 (26-73) |
| HIV viral load | - | - | Undetectable<br>(<50) | Undetectable<br>(<50) |
| Median CD4 count (range) | - | - | 602 (22-1360) | 560(200-1229) |
| Median CD4:CD8 ratio<br>(range) | - | - | 0.74<br>(0.13-3.05) | 0.98<br>(0.37-2.55) |
| **Comorbidities** |  |  |  |  |
| Diabetes, n | - | 1 | 5 | 3 |
| Hypertension/CVD, n | - | 3 | 3 | 6 |
| Renal disease, n | - | - | 2 | 1 |
| Liver disease, n | - | - | 2 | 2 |
| Respiratory disease, n | - | 1 | 3 | 1 |
| Weakened immune system<br>inc. cancer/transplant, n | - | - | 8 | 1 |
| Advanced HIV/HepB co-infection, n | - | - | 4 | - |
| Other | - | - | Splenectomy<br>Sarcoidosis | Splenectomy |
| **Timepoints** |  |  |  |  |
| **Post first dose** |  |  |  |  |
| N= | 17 | 28 | 31 | 32 |
| Median days post-previous<br>dose (range) | 14(12-74) | 19 (12-60) | 20(12-102) | 22(13-82) |
| Vaccine<br>(AZ \| Moderna \| Pfizer) | 3 \| 2 \| 12 | 3 \| 1 \| 24 | 19 \| 2 \| 10 | 21 \| 0 \| 11 |
| **Post second dose** |  |  |  |  |
| N= | 18 | 25 | 30 | 24 |
| Median days post-previous<br>dose (range) | 39(23-67) | 26(15-68) | 20(7-48) | 21(9-52) |
| Vaccine<br>(AZ \| Moderna \| Pfizer) | 3 \| 3 \| 12 | 2 \| 1 \| 23 | 17 \| 1 \| 12 | 12 \| 0 \| 12 |
| **Pre third dose** |  |  |  |  |
| N= | 21 | 26 | 39 | 16 |
| Median days post-previous<br>dose (range) | 129(75-258) | 129(76-236) | 125(72-218) | 119(86-317) |
| Vaccine<br>(AZ \| Moderna \| Pfizer) | - | - | - | - |
| **Post third dose** |  |  |  |  |
| N= | 14 | 25 | 34 | 17 |
| Median days post-previous<br>dose (range) | 21(13-43) | 43(18-129) | 40 (9-149) | 65(7-140) |
| Vaccine<br>(AZ \| Moderna \| Pfizer) | 0 \| 3 \| 12 | 0 \| 2 \| 23 | 1 \| 2 \| 32 | 1 \| 0 \| 16 |

**Figure 1:**
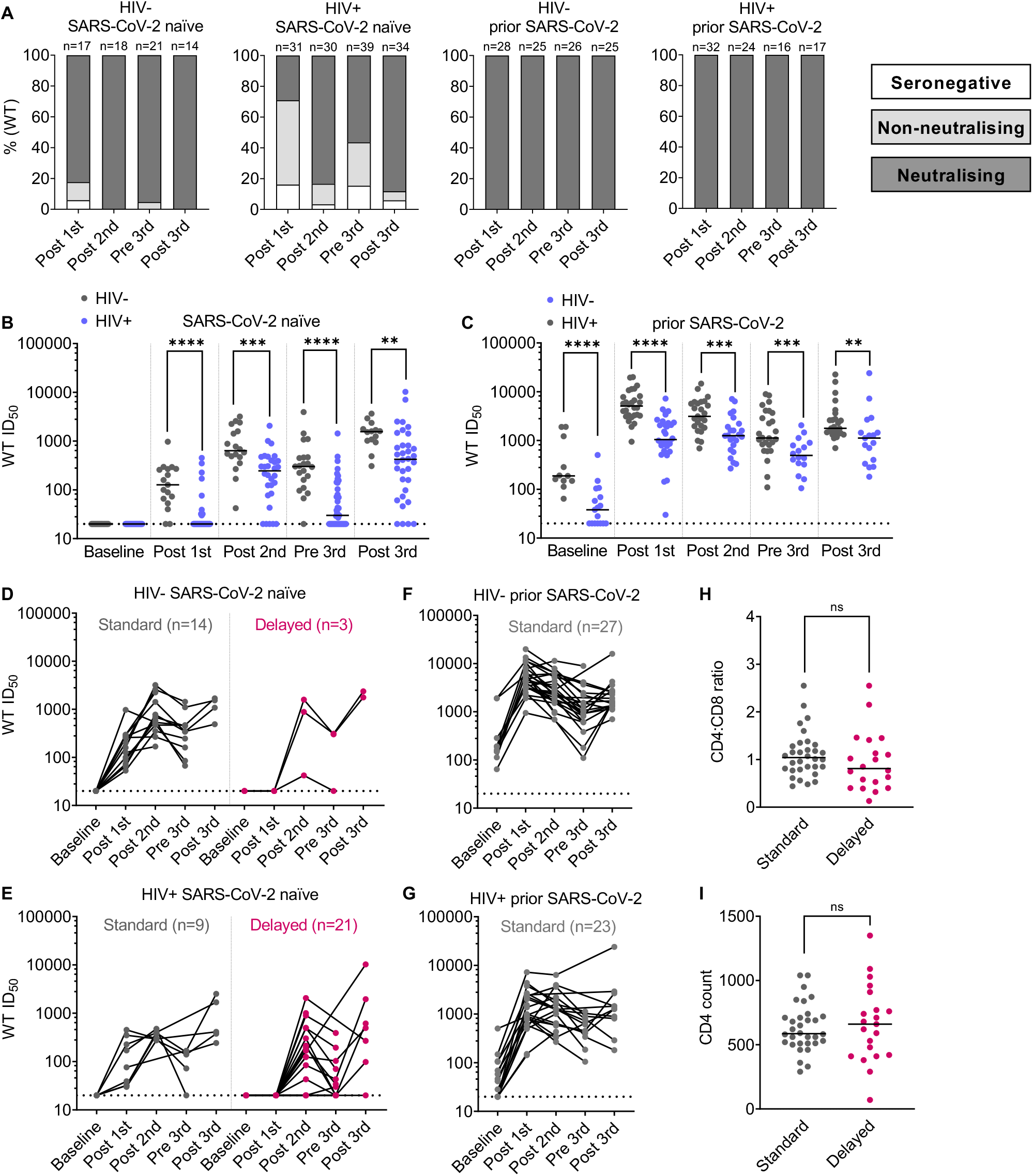
Weaker post vaccination antibody responses in SARS-CoV-2 naïve PLWH. **(A)** Percentage of individuals with detectable neutralizing antibody response, non-neutralizing but binding response, or seronegative at each timepoint as color-coded in the key. The headings above each graph show HIV status and previous SARS-CoV-2 exposure. N numbers for each group are indicated above each column. **(B)** WT pseudovirus neutralization reciprocal 50% inhibitory titers (ID_50_) in PLWH (blue) compared to HIV-negative controls (grey) stratified by vaccination timepoint (on the x-axis) for individuals without prior SARS-CoV-2 infection. The dotted line represents the lower limit of the assay (ID_50_=1:20). Where no neutralization was detected, samples were assigned an ID_50_ of <1:20 as this was the limit of assay detection. Each data point represents the mean of n=2 biological repeats, each measured in duplicates. N numbers match those in (A), Statistical test: Mann Whitney U-test (MWU). **(C)** Shows the equivalent data for those with prior SARS-CoV-2 infection, N numbers match those in (A). **(D)** Longitudinal ID_50_ titers for HIV-negative controls without prior SARS-CoV-2 infection who provided at least two longitudinal samples, including a post first dose sample. Samples that were neutralizing after the first dose are categorised as exhibiting a standard neutralizing response and colored grey, those that only achieve neutralization after the second dose, exhibit a delayed neutralizing response and are color-coded in magenta. N numbers for each category are indicated on the graph. **(E)** Shows the equivalent data for PLWH without prior SARS-CoV-2 infection. **(F)** Shows the equivalent data for HIV-negative controls with prior SARS-CoV-2 infection. **(G)** Shows the equivalent data for PLWH with prior SARS-CoV-2 infection. **(H)** CD4 T cell counts and **(I)** CD4:CD8 T cell ratio for PLWH stratified by standard (grey) or delayed neutralization (magenta). N numbers are as per D-G. Statistical test: MWU.

Notably, PLWH had lower titers of nAbs than HIV-negative controls at all timepoints regardless of prior SARS-CoV-2 infection **(Figure 1B, C)**. Overall, a similar trend was seen in binding responses **(Supplementary Figure 1A, B)**, and nAb titers correlated significantly with both binding titers for S1 IgG and nAb titers obtained from a live virus neutralization assay **(Supplementary Figure 1C, D)**, as previously reported (Brouwer et al., 2020; Graham et al., 2021). Given that this observational cohort includes a mixture of SARS-CoV-2 vaccine types, it was notable that both binding and neutralizing titers remained significantly lower in PLWH compared to controls when only those who had received mRNA-based vaccines were considered **(Supplementary Figure 1E, F)**. A similar analysis for viral vector-based vaccines was not feasible due to insufficient numbers in the control group. Both at the pre- and post-third vaccine dose timepoints, there were more SARS-CoV-2 naïve PLWH that fail to produce nAbs **(Figure 1A, 1B, Supplementary Figure 1A)** compared to the control group. This could be biased by the cross-sectional nature of the analysis as at the pre-third vaccine dose timepoint, additional PLWH were recruited, some with complex co-morbidities. However, the observed differences persisted when PLWH were stratified for co-morbidities **(Supplementary Figure 1G)**.

Longitudinal samples from 53 PLWH and 44 controls were then evaluated to assess binding antibody responses and nAbs over time after each vaccine dose. These included samples after the first dose and for at least one additional timepoint, often including a baseline, post-second, pre-third and post-third sample **(Figure 1D)**. This analysis revealed two clear trajectories of the development of neutralization, firstly where nAbs were detected after a single vaccine dose (Gilbert et al., 2022), defined here as “standard neutralization”, and secondly where neutralization was not achieved until after the second dose or later, defined as “delayed neutralization”. Most HIV-negative controls without prior SARS-CoV-2 infection show a standard neutralization profile, with only 3 individuals failing to mount a neutralizing response until after the second dose **(Figure 1D)**, and a similar effect was seen with binding responses **(Supplementary Figure 1H-K)**. In contrast, two-thirds of SARS-CoV-2 naïve PLWH did not make a detectable neutralizing response until after the second dose and a substantial proportion of them lost detectable neutralizing activity before the third dose **(Figure 1A, E)**. However, both PLWH and HIV-negative controls with a history of SARS-CoV-2 infection made a standard neutralizing response **(Figure 1F, G)**. Therefore, having identified this delayed neutralization phenotype in SARS-CoV-2 naïve PLWH, we have evaluated its relationship with total CD4 T cell counts, which are known to be important for SARS-CoV-2 vaccine responses in PLWH (Hassold et al., 2022; Nault et al., 2021; Noe et al., 2021; Touizer et al., 2021). No significant difference was seen in median CD4 T cell count or CD4:CD8 T cell ratio between PLWH with standard or delayed neutralization profiles **(Figure 1H, I);** or correlate either with the rapid development of neutralization **(Supplementary Figure 1M, N)**.

### Delayed neutralization is associated with lower frequency of spike-specific MBCs and a perturbed MBC global phenotype

Spike is the SARS-CoV-2 glycoprotein and is the sole antigen in most vaccines. It has been previously shown that infection and vaccination produce spike-specific MBCs in proportion to serological responses (Cohen et al., 2021; Dan et al., 2020; Goel et al., 2021; Jeffery-Smith et al., 2022; Terreri et al., 2022). Given that the delay in neutralization observed more frequently in PLWH was not clearly associated with peripheral CD4 T cell counts, we next assessed the relationship with global MBCs and spike-reactive MBC frequency and phenotype, using a previously validated flow cytometry panel, with memory B cells defined as CD19+ CD20+ CD38^lo/-^ IgD-**(Supplementary Figure 2)**. This analysis was performed on available PBMC samples after the first vaccine dose, using SARS-CoV-2 naïve baseline samples to determine the antigen-specific gate **(Figure 2A)**. We observed a significantly lower frequency of spike-specific MBCs in SARS-CoV-2 naïve participants after the first dose as compared to those with a history of prior infection, regardless of HIV status **(Figure 2B)**. Moreover, a lower frequency of spike-specific MBCs was observed in SARS-CoV-2 naïve participants who had a delayed neutralization response, although notably there was a small number of donors in the standard neutralization group **(Figure 2C)**. In line with this, the percentage of spike-specific MBCs showed a strong correlation with the nAb titer **(Figure 2D)** in agreement with previous findings during SARS-CoV-2 convalescence (Jeffery-Smith et al., 2022).

**Figure 2:**
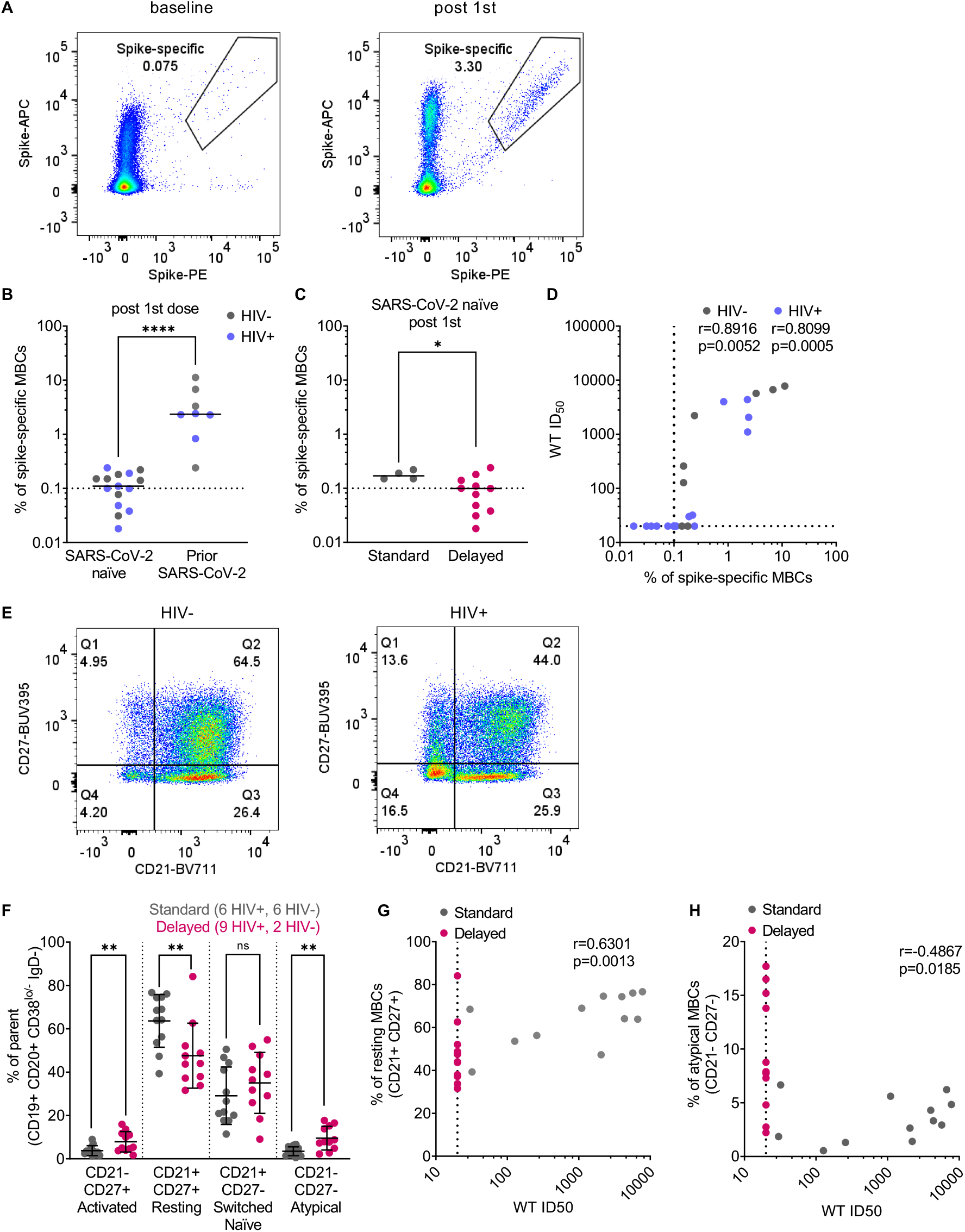
Neutralization titer is associated with the frequency of spike-specific MBCs after the first vaccine dose. **(A)** Spike-specific MBCs (CD19+ CD20+ CD38^lo/mid^ IgD-excluding switched naïve CD27-CD21+ cell) according to spike-PE and spike-APC in a representative naïve pre-vaccine sample (left) or representative post-vaccine sample (right) after the first vaccine dose. **(B)** Percentage of spike-specific MBC after the first vaccine dose stratified by prior SARS-CoV-2 infection, statistical test: M-Whitney U test (MWU). Dotted lines represent lower limit of sensitivity of the assay (0.1% spike-specific MBCs, based on (Jeffery-Smith et al., 2022). **(C)** Percentage of spike-specific MBCs in SARS-CoV-2 naïve donors after the first vaccine dose, stratified by delayed (magenta) or standard (grey) neutralization profile, statistical test: MWU. Dotted lines represent lower limit of sensitivity of the assay (0.1% spike-specific MBCs) **(D)** Correlation of the percent of spike-specific MBC with WT ID_50_ titers stratified by PLWH (blue) and controls (grey) after the first dose, statistical test: Spearman’s rank correlation coefficient. **(E)** Distribution of MBCs (CD19+ CD20+ CD38^lo/mid^ IgD-) subtypes according to CD27-BUV395 and CD21-BV711 in a representative HIV-negative donor sample (left) or PLWH donor sample (right). **(F)** Percentage of MBC subtypes (activated CD27+ CD21-; resting CD27+ CD21+; switched naïve; switched naïve CD27-CD21+ and CD27-CD21-atypical) after the first vaccine dose stratified by delayed or standard neutralization profile. Statistical test: MWU **(G)** Correlation of the percentage of resting CD27+ CD21+ MBCs with WT ID_50_ titers stratified by delayed (magenta) or standard (grey) neutralization profile after the first vaccine dose, statistical test: Spearman’s rank correlation coefficient. **(H)** Correlation of the percent of switched naïve CD27-CD21+ MBCs with WT ID_50_ titers stratified by delayed (magenta) or standard (grey) neutralization profile after the first vaccine dose, statistical test: Spearman’s rank correlation coefficient.

Subsequent gating on CD21 and CD27 expression allowed the identification of four populations of class-switched MBCs: CD21-CD27-atypical MBCs (also known as tissue-like memory); CD21-CD27+ activated MBCs; CD21+CD27+ classical resting MBCs and CD21+ CD27-switched naïve (also known as intermediate memory) MBCs **(Figure 2E)** as previously described (Jeffery-Smith et al., 2022). Global defects in the balance of these MBC subsets have been identified previously in PLWH (reviewed in (Moir and Fauci, 2017)), including those on ART (Pensieroso et al., 2013), with increased numbers of activated and atypical MBCs concurrent with a decrease in resting MBCs. This phenotype is exemplified in (**Figure 2E**) for a PLWH and a HIV-negative control. We have hypothesised that these inherent defects may have an impact on the quality of serological responses after SARS-CoV-2 vaccination. Global phenotyping of the MBC response after the first vaccine dose revealed that individuals with delayed neutralization, consisting largely of PLWH, had significantly lower numbers of resting MBCs (CD21+ CD27+) and greater numbers of both CD21-CD27+ activated MBCs and CD21-CD27-atypical MBCs compared to those with standard neutralization **(Figure 2F)**. Moreover, lower frequencies of resting MBCs correlated with lower nAb titers **(Figure 2G)**. Higher levels of atypical MBCs significantly correlated with lower nAb titers, although the strength of this association was relatively weak (r=-0.4867) **(Figure 2H)**. Together these findings suggest that the MBC subset perturbations seen in PLWH could account for the lower serological output.

### Improved neutralization breadth after the third SARS-CoV-2 dose in PLWH but lower nAb responses persist and are associated with global, but not spike-specific, MBC dysfunction

To assess the breadth of nAb responses across the cohort, samples from all timepoints were tested against an Omicron pseudovirus (BA.1 strain), which represented the dominant circulating strain at the time of the post third vaccine dose sampling. Due to the substantial antigenic changes in the Omicron spike (McCallum et al., 2022), in participants with no prior infection, over 50% of HIV-negative controls and more than 90% of PLWH were not able to neutralize Omicron after the first vaccine dose **(Figure 3A)**. The second dose enabled most of the control group to mount a neutralizing response whereas only a quarter of SARS-CoV-2 naïve PLWH had nAbs against Omicron. In the SARS-CoV-2 naïve groups, the third dose enabled 100% of HIV-negative controls to neutralize Omicron and increased the frequency of neutralization among PLWH to over 70% **(Figure 3A)**. As in the analysis of WT neutralization for individuals without prior SARS-CoV-2 infection, median Omicron ID_50_ titers were lower in SARS-CoV-2 naïve PLWH compared to HIV-negative controls at all timepoints **(Figure 3B)**. Additionally, there was no significant difference when individuals with complex co-morbidities were removed from the PLWH cohort at the third vaccine dose **(Supplementary Figure 3C)** or whether they had previously been infected with SARS-CoV-2. These data suggest that the third vaccine dose was effective in both boosting nAb titer and broadening the response to Omicron, especially in SARS-CoV-2 naïve PLWH, thus rendering their responses closer to those of SARS-CoV-2 naive HIV-negative controls **(Figure 3B-C)**.

**Figure 3:**
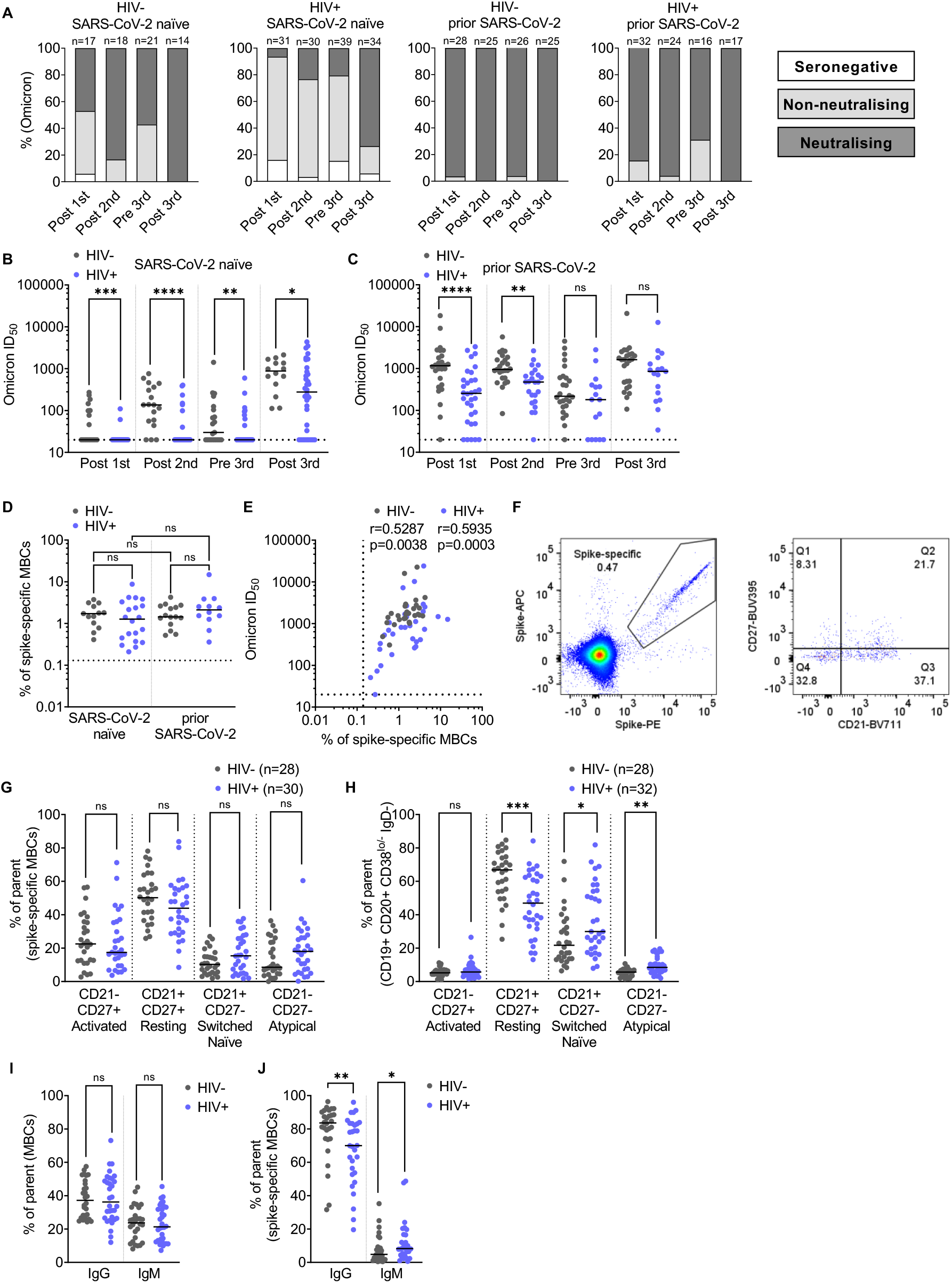
Improved neutralization against Omicron after the third vaccine dose in PLWH accompanied by minimal alteration in the spike-specific MBC phenotype. **(A)** Percentage of individuals with detectable neutralizing response, non-neutralizing but binding response, or seronegative at each timepoint as color-coded in the key (neutralization against Omicron pseudovirus). Headings above each graph show the HIV status and previous SARS-CoV-2 exposure. N numbers for each group are indicated above each column. **(B)** Omicron pseudovirus neutralization ID_50_ in PLWH (blue) compared to HIV-negative controls (grey) stratified by vaccination timepoint (on the x-axis) for individuals without prior SARS-CoV-2 infection. Statistical test: Mann-Whitney U test (MWU). **(C)** Shows the equivalent data for those with prior SARS-CoV-2 infection, N numbers match those in (A). **(D)** Percentage of spike-specific MBCs in PLWH (blue) and HIV-negative donors (grey) after the third vaccine dose stratified by SARS-CoV-2 infection. Statistical test: MWU. **(E)** Correlation between Omicron ID_50_ titers and percentage of spike-specific MBCs in PLWH (blue) and HIV-negative donors (grey) after the third vaccine dose. Statistical test: Spearman’s rank correlation coefficient. **(F)** Representative gating strategy to identify spike-specific MBCs subtypes. **(G)** Percentage of spike-specific MBCs subtypes (activated CD27+ CD21-; resting CD27+ CD21+; switched naïve; switched naïve CD27-CD21+ and CD27-CD21-atypical) after the third vaccine dose in PLWH (blue) and HIV-negative donors (grey). Statistical test: MWU. **(H)** Percentage of MBCs subtypes (activated CD27+ CD21-; resting CD27+ CD21+; switched naïve; switched naïve CD27-CD21+ and CD27-CD21-atypical) after the third vaccine dose in PLWH (blue) and HIV-negative donors (grey). Statistical test: MWU. **(I)** Percentage of IgG and IgM in MBCs (excluding switched naïve CD27-CD21+ fraction) after the third vaccine dose in PLWH (blue) and HIV-negative donors (grey). Statistical test: MWU. **(J)** Percentage of IgG and IgM in spike-specific MBCs after the third vaccine dose in PLWH (blue) and HIV-negative donors (grey). Statistical test: MWU.

Next, we evaluated cross-sectionally the B cell phenotype after the third vaccine dose. In contrast to the first vaccine dose, there was no significant difference between the frequency of spike-specific MBCs when individuals were stratified by whether they had been previously infected with SARS-CoV-2 or not **(Figure 3D)** regardless of HIV status. However, the frequency of spike-specific MBCs after the third dose correlated with Omicron titers **(Figure 3E)**. This suggests that after three vaccine doses these individuals had mounted a specific B cell response, and that the quantity of spike-specific B cells remained linked to the improved neutralization potency and breadth observed **(Figure 3A-C)**. Given that all individuals assessed after the third dose made a robust spike-specific MBC response, we wanted to evaluate further whether alterations in spike-specific MBC phenotype also contributed to differences in serum neutralization **(Figure 3A, B, F, Supplementary Figure 3A-B)**. Spike-specific B cells were found to be comparable across the different MBC subsets in both PLWH and HIV-negative controls, except for a trend to fewer spike-specific resting MBCs in PLWH as compared to controls **(Figure 3G)**. This was the case even though the global MBC population for these post third vaccine dose samples showed classical anomalies in MBCs associated with HIV infection **(Figure 3H)**. These data suggest that SARS-CoV-2 serum antibody responses are lower potentially because of a global MBC disturbance thereby limiting the overall B cell response. In line with this proposal, we anticipated that underlying global MBC disturbances would also influence the efficiency of the antigen-specific B cell response in other ways, beyond limiting the number of spike-specific MBCs, for example by limiting class-switching. Indeed, this is supported by our data showing similar levels of IgG+ and IgM+ global MBCs in both groups **(Figure 3I)** but a significantly lower level of spike-specific IgG+ MBCs in PLWH after the third vaccine dose as compared to controls, and conversely a higher frequency of spike-specific IgM+ MBCs **(Figure 3J)**.

### SARS-CoV-2 vaccination induces robust T cell responses that cross-recognize variants in PLWH

To increase our understanding of the complementary role of cellular immunity after vaccination, we have examined T cell responses in our cohort, including their reactivity to SARS-CoV-2 variants. The magnitude of spike-specific T cell responses was assessed cross-sectionally by IFN-γ-ELISpot using overlapping peptide (OLP) pools covering the complete sequences of the WT spike glycoprotein as previously described (Alrubayyi et al., 2021). The majority of PLWH had detectable SARS-CoV-2-specific T cell responses at levels comparable to HIV-negative individuals following each vaccine dose (**Figure 4A-C)**. A greater magnitude of spike-specific T cells was observed in individuals with prior SARS-CoV-2 infection, irrespective of HIV status (**Figure 4A-C**) in keeping with previous reports (Lozano-Ojalvo et al., 2021; Prendecki et al., 2021; Reynolds et al., 2021). There were no detectable T cell responses in a small number of PLWH with no prior exposure to SARS-CoV-2 across all timepoints. These were participants with incomplete immune reconstitution on ART and/or additional co-morbidities, such as transplant recipients on immunosuppressive therapy (**Figure 4A-C**). Next, we examined the longitudinal evolution of T cell responses in a subgroup of donors with available PBMC samples. In SARS-CoV-2 naïve individuals, spike-specific T cell responses increased following the first vaccine dose, peaked after the second dose and were maintained after the third vaccine dose (**Figure 4D)**. In one HIV-positive, SARS-CoV-2-naïve donor with advanced immunosuppression and persistently low CD4 T cell count of 100 cells/μL on ART, a third dose (mRNA) vaccine was able to elicit a T cell response despite no evidence of neutralization (**Figure 4D**). A higher proportion of PLWH without prior SARS-CoV-2 infection had detectable T cell responses at baseline compared to HIV-negative controls, which could represent the presence of cross-reactive responses to other pathogens, probably to related coronaviruses (**Figure 4D**) (Braun et al., 2020; Grifoni et al., 2020; Le Bert et al., 2020; Mateus et al., 2020; Sekine et al., 2020). However, due to the small number of participants with detectable T cell responses at baseline, this study was not powered to detect any association between the presence of cross-reactive T cells and magnitude of vaccine-induced T cell responses. In donors with prior SARS-CoV-2 infection, there was a boosting effect to spike-specific T cells following the first vaccine dose in both study groups (**Figure 4D**). In parallel we have tested T cell responses to CMV-pp65 and HIV-gag peptide stimulation within the same individuals across all timepoints. Overall, PLWH with no prior exposure to SARS-CoV-2 had robust responses to CMV-pp65 stimulation, as expected given their higher CMV seroprevalence compared to HIV negative donors. CMV-specific responses in these individuals were higher compared to SARS-CoV-2 and Gag-specific responses following each vaccine dose (**Supplementary Figure. 4 A-C)**. Prior SARS-CoV-2 exposure resulted in comparable SARS-CoV-2 and CMV-pp65 T cell responses after the third vaccine dose in PLWH (**Supplementary Figure. 4C)**. No significant differences were detected between SARS-CoV-2 and CMV-specific responses in HIV-negative individuals (**Supplementary Figure. 4A-C)**. Overall, these results demonstrate a robust induction of T cell responses to SARS-CoV-2 vaccination in PLWH despite attenuated antibody responses.

**Figure 4:**
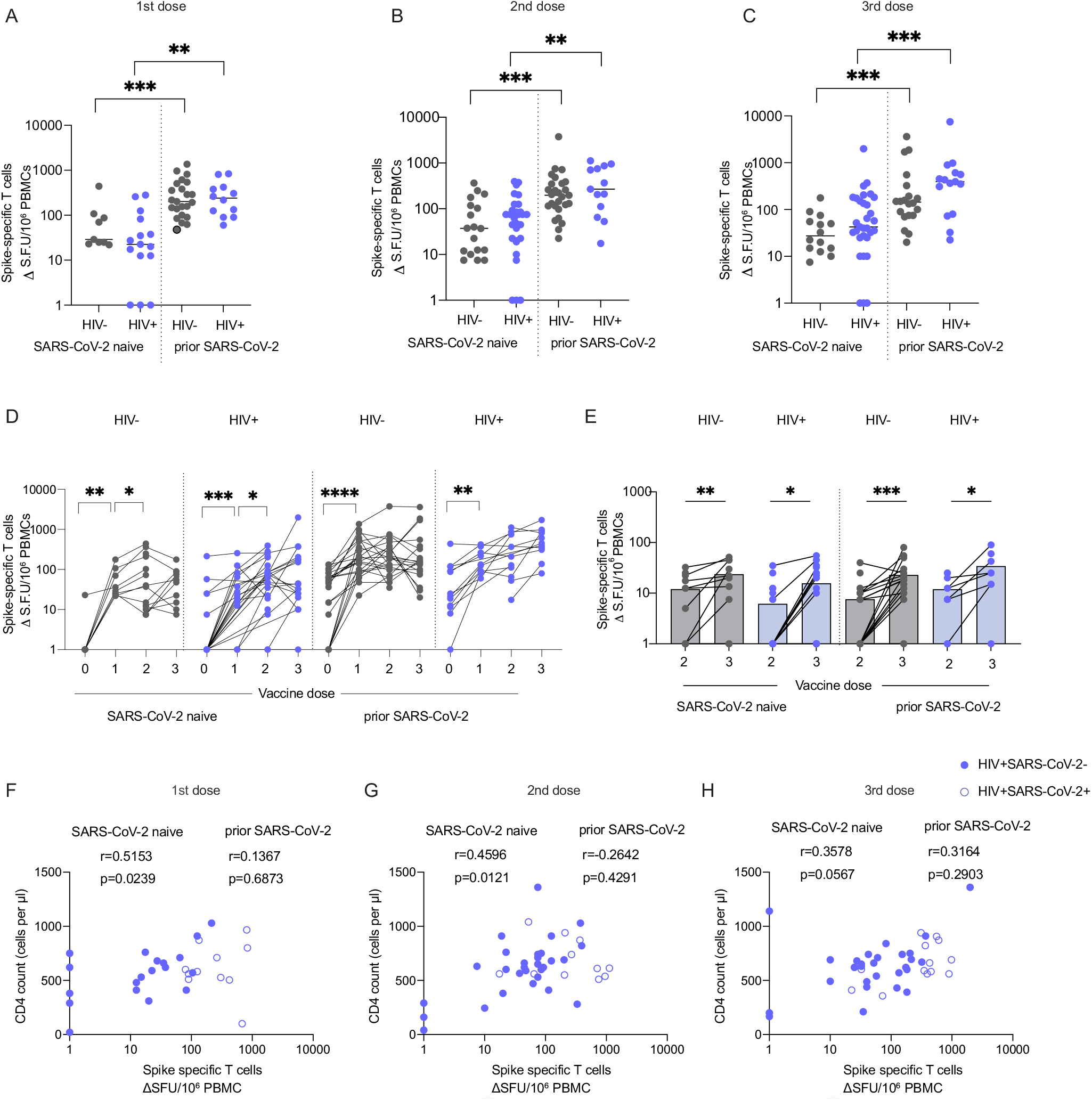
Comparable magnitude of spike-specific T-cell responses following SARS-CoV-2 vaccination in HIV-positive and HIV-negative individuals. **(A-C)** Cross-sectional analysis of the magnitude of the IFN-γ-ELISpot responses to the SARS-CoV-2 spike peptide pools in HIV-negative (grey) and HIV-positive (blue) individuals, with or without prior SARS-CoV-2 infection following first dose (A) second dose (B) and third dose (C). (HIV-SARS-CoV-2-first dose n=9, second dose n=18, third dose n=14; HIV+SARS-CoV-2-frist dose n=15, second dose n=29, third dose n=31; HIV-SARS-CoV-2+ first dose n=23, second dose n=27, third dose n=20; HIV+SARS-CoV-2+ first dose n=12, second dose n=13, third dose n=15). Statistical test: Mann-Whitney U-test (MWU). **(D)** Longitudinal analysis of the spike specific T cell responses in PLWH and HIV-negative subjects. Statistical test: Wilcoxon matched-pairs sign rank test (WMP). **(E)** Longitudinal and cross-sectional analysis of the magnitude of T cell responses to B.1.1.529 after two or three vaccine doses (n=11 HIV-SARS-CoV-2-, n=20 HIV+SARS-CoV-2-, n=22 HIV-SARS-CoV-2+, n=10 HIV+SARS-CoV-2+). Statistical test: MWU and WMP. **(F-H)** Correlation between the CD4 T cell count in HIV-positive individuals and magnitude of spike-specific T cell responses after first dose (F), second dose (G), and (H) third dose. Statistical test: Spearman’s rank correlation coefficient

Previous work has demonstrated that T cell responses are largely retained against variants of concern (VOCs), including the highly transmissible BA.1 Omicron variant, and therefore may be important when antibody levels wane or new variants emerge that can partly escape antibody responses. To determine T cell reactivity to VOCs, we assessed T cell responses to the mutated regions, including Omicron, in our study cohort. The magnitude of T cell responses against B.1.1.529 was comparable between PLWH and HIV-negative donors regardless of prior SARS-CoV-2 infection **(Figure 4E)**. Notably, responses were further enhanced by a third vaccine dose in all donors, irrespective of prior SARS-CoV-2 infection or HIV status and in keeping with the beneficial effect of a third vaccine dose in boosting humoral responses **(Figure 4E)**. T cell reactivity to Omicron and other VOCs, including Alpha, Beta and Delta, was comparable between HIV-negative and PLWH with or without prior SARS-CoV-2 infection after three vaccine doses, and these responses were maintained against the ancestral Wuhan Hu-1 spike peptide pool, reinforcing the relative resilience of T cell responses to spike variation (**Supplementary Figure 4D-E)**. We noted that three HIV-negative and five HIV-positive individuals, regardless of prior SARS-CoV-2 infection, had no detectable T cell responses to the Wuhan Hu-1 peptide pool, covering only the affected regions of spike. This could be in part due to the VOC mutations occurring in regions that are poorly targeted by T cell responses in some individuals (Reynolds et al., 2021).

Although spike-specific T cell responses were detected at similar frequencies across all groups **(Figure 4A-C)**, there was variation in the magnitude of responses. To better understand the factors underlying this heterogeneity, we examined the role of various HIV parameters (Alrubayyi et al., 2021). We have previously reported an association between the CD4:CD8 T cell ratio and total SARS-CoV-2 responses, especially against the nucleocapsid (N) and membrane (M) protein, in PLWH recovering from COVID-19 disease (Alrubayyi et al., 2021). No correlation was observed between the CD4:CD8 T cell ratio and spike-specific T cell responses following vaccination in our cohort (**Supplementary Figure 4F-H)**. However, a positive correlation was detected between the CD4 T cell count and spike-specific T cell responses after the first vaccine dose (r=0.5153) in SARS-CoV-2 naïve PLWH (**Figure 4E**). This association was weaker after the second vaccine dose (r=0.4596) and non-significant after the third dose (**Figure 4F, G**). Together these observations suggest that an effective helper T cell response could drive the induction of cellular immunity following vaccination in individuals without prior exposure to SARS-CoV-2. However, the lack of an association between CD4 T cell counts and antibody responses further underlines the relative importance of HIV-associated B cell defects in modulating the induction of effective humoral immunity in addition to potential insufficient T cell priming.

### A proportion of PLWH had low or absent nAbs (ID_50_ <150) but detectable T cell responses following vaccination

We examined next the relationship between humoral and cellular responses by comparing antibody responses and neutralization titers with T cell responses detected by IFN-γ-ELISpot following SARS-CoV-2 vaccination. Overall, spike-specific T cells following the first, second and third vaccine doses correlated positively with respective nAb titers in HIV-negative and PLWH. These associations were stronger in PLWH after the first (r=0.5402; p=0.0014) and second dose of vaccine (r=0.5038, p=0.0004), similarly to HIV-negative controls **(Figure 5A-C)**. Similar associations were observed for S1 IgG binding titers (**Supplementary Figure 5A-C)**. One HIV-positive SARS-CoV-2 naïve donor with a low CD4 T cell count of 40 cells/μL on ART, and one individual with relapsed lymphoma, both had no detectable humoral and cellular responses after 2 or 3 doses of mRNA vaccine. Interestingly a proportion of PLWH, in particular those without prior SARS-CoV-2 infection, had low or absent nAbs (ID_50_ <150) but detectable T cell responses following vaccination (**Figure 5A-C**). To better visualise these relationships in SARS-CoV-2 naïve individuals, we ranked T cell responses after second and third doses according to the magnitude of neutralizing antibodies (**Figure 5E-G**). All of the HIV-negative donors had detectable cellular and neutralizing antibodies (**Figure 5D**). However, a proportion of SARS-CoV-2 naïve PLWH with low or absent nAbs (n=9 out of 10) had measurable cellular responses to the spike protein after two vaccine doses **(Figure 5E)**. These donors were all controlled on ART with a median CD4 T cell count of 680 cells/μL and no significant underlying co-morbidity (Supplementary Table 1). Although all HIV-negative individuals had both detectable nAbs and cellular responses post third dose (**Figure 2F**), a small number of PLWH SARS-CoV-2 naïve donors (n=7 out of 9) had detectable T cell responses in the absence of, or only low-level, neutralization **(Figure 5G)**. Similarly, these donors were all well controlled on ART with a median CD4 T cell count of 492 cells/μL. One of these donors who presented with advanced HIV infection had a persistently low CD4 T cell count (100 cells/μL), and one of the donors recruited after a third vaccine dose had a previous splenectomy. These data suggest that in a small proportion of PLWH, serological non-responders or with evidence of low-level neutralization, cellular immune responses may play an important compensatory role.

**Figure 5:**
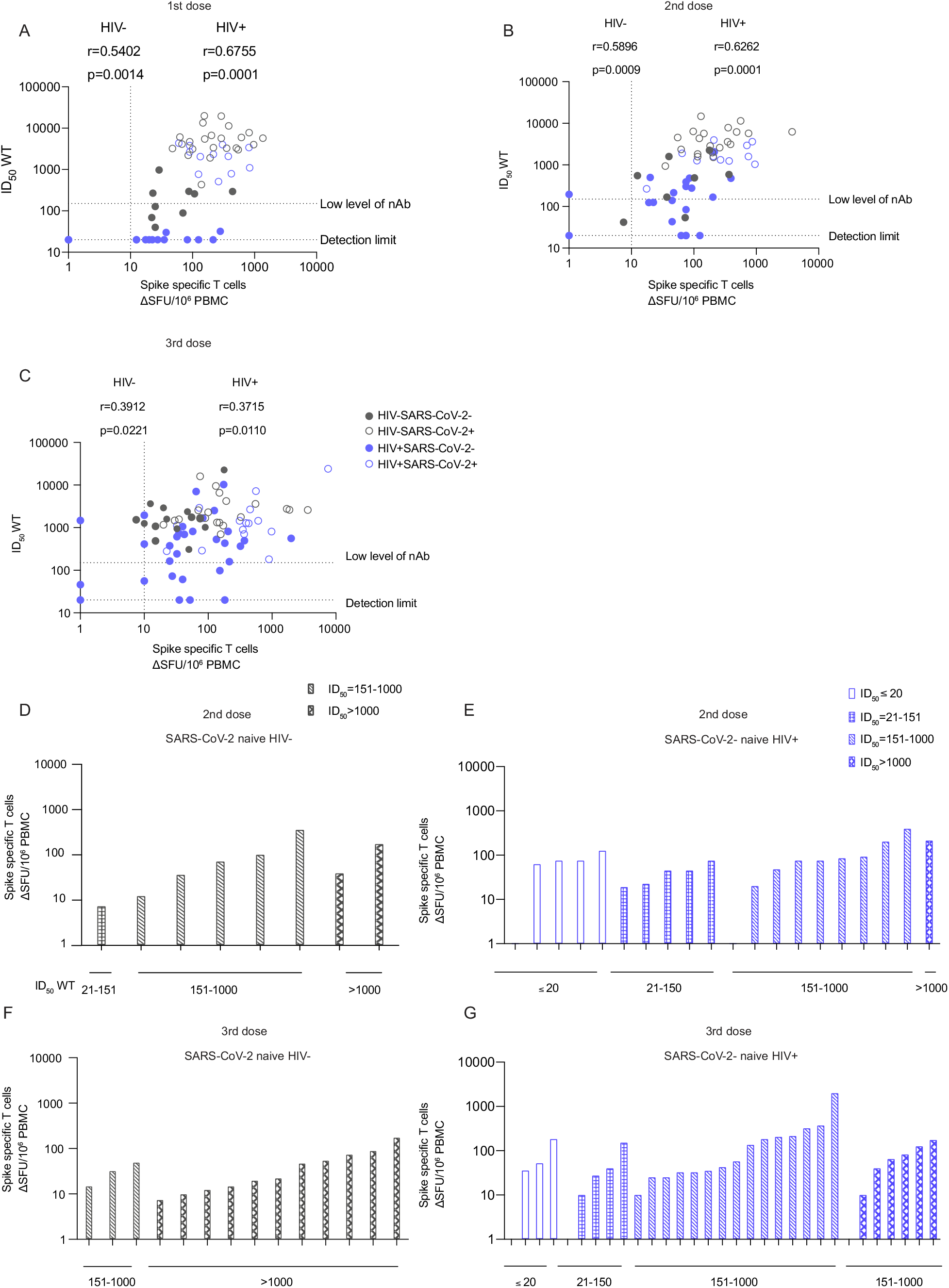
Interrelations between humoral and cellular responses following SARS-CoV-2 vaccination in HIV positive and HIV negative individuals. **(A-C)** Correlation of spike-specific T cell responses with nAb titers after first dose (A) second dose (B) and third vaccine dose (C) of vaccine in HIV-negative and HIV-positive donors, with or without prior SARS-CoV-2 infection (limit of detection ID_50_=20, low level of nAb ID_50_=150). Statistical test: Spearman’s rank correlation coefficient. **(D-E)** Hierarchy of the spike-specific T cell responses ordered by their nAb titers in HIV-negative (D) and HIV-positive (E) SARS-CoV-2 naïve donors after two vaccine doses. **(F-J)** Hierarchy of the spike-specific T cell responses after three vaccine doses in HIVnegative (F) and positive (J) SARS-CoV-2 naïve participants.

### PLWH with suboptimal serological responses demonstrate an expansion of CXCR3^+^CD127^+^ CD8^+^ T cells after two doses of SARS-CoV-2 vaccination

The presence of detectable T cell responses in a subgroup of SARS-CoV-2 naïve HIV-positive donors with low or absent nAbs after two or three vaccine doses prompted us to further evaluate the phenotype of the T cell compartment. We have compared T cell immune signatures in SARS-CoV-2 naïve PLWH with potent neutralization titers (>1:150) and functional T cell responses (PLWH SARS-CoV-2 naive nAb^high^T^+^, n=9), with SARS-CoV-2 naïve PLWH with low/absent nAbs and a functional T cell responses (PLWH SARS-CoV-2-nAb^-/low^T^+^, n=9). Both groups were age and sex matched, well controlled on ART and with a similar median CD4 T cell count **(Supplementary Table 1**). We have used an unbiased approach and unsupervised high-dimensional analysis, global t-distributed stochastic neighbour embedding (t-SNE), followed by FlowSOM clustering, in circulating T cell populations in the two groups. Ten major CD4 and CD8 T cell subsets were examined using a combination of various activation and differentiation markers, including CD45RA, CCR7, CD127, CD25, CXCR3, CXCR5, PD-1, and CD38 (**Figure 6A and Supplementary Figure 6A-B**). There was no difference in the frequencies of the main T cell subsets in the two groups (**Supplementary Figure 6D**). Among CD4 T cells, there was a reduction in circulating CXCR3^+^CXCR5^+^ T follicular helper (T_FH_) subsets observed in HIV-positive nAb^-/low^ compared to nAb^+^ donors (**Figure 6 A,B)**. The reduced abundance of CXCR3^+^CXCR5^+^ T_FH_ in nAb^-/low^ HIV-positive subjects was further confirmed by manual gating (**Figure 6C, D and Supplementary 3C)**. CXCR3^+^CXCR5^+^ T_FH_ cells correlated with SARS-CoV-2 neutralization levels in HIV-positive SARS-CoV-2 naïve individuals (r=0.5294 p=0.02388) (**Figure 6E)**, suggesting that reduced availability of T_FH_ cells could influence the magnitude of vaccine-induced SARS-CoV-2 antibody responses.

**Figure 6:**
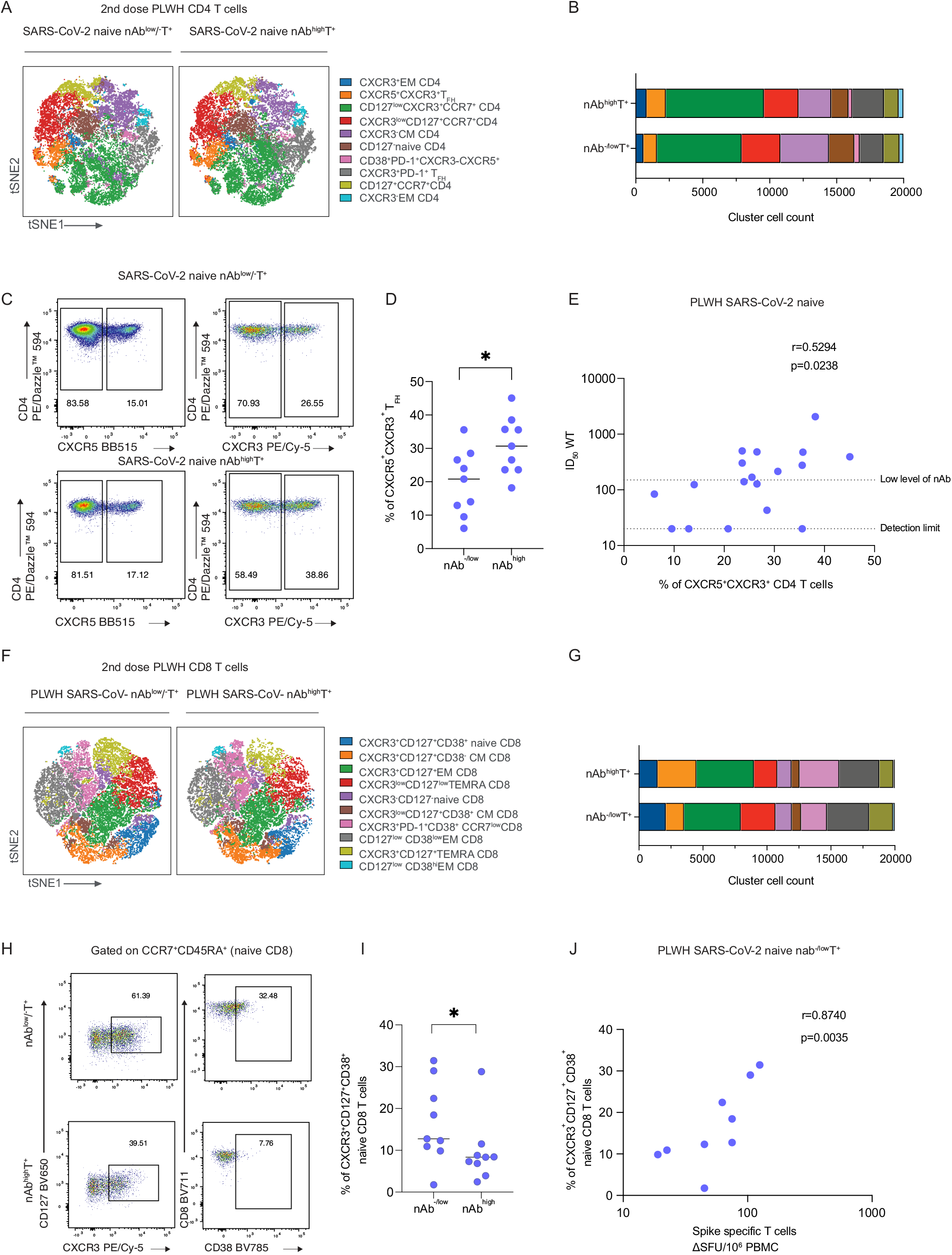
Phenotypic characterization of CD4 and CD8 T cells from SARS-CoV-2 naïve HIV positive individuals according to their neutralization levels. **(A)** viSNE map of FlowSOM metaclusters of CD4 T cells from HIV positive SARS-CoV-2 naïve subjects after two vaccine doses (nab^-/low^= no neutralization or low level of neutralization, nAb^high^=high neutralization level; n=9 in each group). Each point on the high-dimensional mapping represents an individual cell, and metaclusters are color-coded. **(B)** Cell count of each FlowSOM metaclusters out of total CD4 T cells (20,000 cells/group). **(C)** Representative flow plots from a nAb^-/low^ and nAb^high^ SARS-CoV-2 naïve HIV-positive donor showing expression of CXCR5 and CXCR3 within CD4 T cells. **(D)** Summary analysis of the percentage of CXCR5^+^CXCR3^+^CD4 T cells (n=9 for each group). Statistical test: Mann-Whitney U-test (MWU). **(E)** Correlation between frequency of CXCR5^+^CXCR3^+^CD4 T cells and ID_50_ neutralization level in nAb^-/low^ and nAb^high^ SARS-CoV-2 naïve HIV-positive individuals after two vaccine doses. Statistical test: Spearman’s rank correlation coefficient. **(F)** viSNE map of FlowSOM metaclusters of CD8 T cells from nAb^-/low^ and nAb^high^ HIV-positive SARS-CoV-2 naïve subjects after two doses of the vaccine (n=9 in each group). **(G)** Cell count of each CD8 FlowSOM metaclusters out of total CD8 T cells (20,000 cells/group). **(H)** Representative flow plots from a nAb^-/low^ and nAb^high^ SARS-CoV-2 naïve HIV-positive donor showing expression of CXCR3, CD127, and CD38 within naïve CD8 T cells. **(I)** Summary analysis of the percentage of CD127^+^CXCR3^+^CD38^+^niave CD8 T cells (n=9 for each group). Statistical test: MWU. **(J)** Correlation between proportion of CD127^+^CXCR3^+^CD38^+^niave CD8 T cells and SARS-CoV-2 specific T cell responses in nAb^-/low^ HIV-positive SARS-CoV-2 naïve subjects. Statistical test: Spearman’s rank correlation coefficient.

We have next examined the CD8 T cell compartment in the two groups. Notably, a prominent cluster delineated by the expression of CXCR3^+^CD127^+^CD38^+^CCR7^+^CD45RA^+^ was significantly enriched in PLWH SARS-CoV-2-nAb^-/low^T^+^ (**Figure 6F, G and Supplementary 6B)**. The higher abundance of CXCR3^+^CD127^+^CD38^+^CCR7^+^CD45RA^+^ cells in PLWH SARS-CoV-2-nAb^-/low^T^+^ was further confirmed by manual gating (p=0.04) (**Figure 6H, I and Supplementary 6C)**. Correlation analysis of these populations showed a positive association between their frequencies and SARS-CoV-2-specific T cell responses following two vaccine doses in PLWH with nAb^-/low^ **(Figure 6)**, supporting the notion that these subsets could contribute to the observed induction of T cell responses in PLWH who lacked or generated low nAb responses. Overall, our analysis of the global T cell profile of individuals with low/absent nAbs but detectable functional T cell responses revealed that reduced availability of T_FH_ cells could contribute to the serological defect observed in conjunction with the previously highlighted imbalance in MBCs. Moreover, we have identified a subset of CD8 T cells that is overrepresented in PLWH with low/absent nAbs and may enable stronger functional T cell responses, supported by recent findings showing that CXCR3^**+**^ CD8 T cells are polyfunctional and associated with survival in critical SARS-CoV-2 patients, and have been observed in other immunosuppressed groups (Adam et al., 2021; Gao et al., 2022).

## DISCUSSION

Accumulating evidence suggests that a broad and well-coordinated immune response is required for protection against severe COVID-19 disease. The emergence of VOCs with increased ability to evade nAbs has reinforced the need for a more comprehensive assessment of adaptive immunity after vaccination, especially in more vulnerable groups including some PLWH. Our data show that PLWH who are well controlled on ART, elicited poorer humoral responses, in terms of magnitude and neutralizing ability compared to HIV-negative donors following first, second and third doses of SARS-CoV-2 vaccine. This was related to global B cell but not antigen-specific B cell dysfunction, thereby providing new insights into what enables a fully-fledged vaccine response. In contrast, T cell responses were comparable in the two groups and detectable, even in a small group of PLWH with very poor serological responses, suggesting a potentially important non-redundant immunological role for functional T cells. Overall, our data reinforce the beneficial effect of an additional vaccine dose in boosting adaptive immune responses (Vergori et al., 2022), especially against circulating VOCs in this patient group.

Weaker humoral responses were observed in PLWH compared to HIV-negative controls after each dose of vaccine when matched by prior SARS-CoV-2 status. While the third dose largely narrowed the gap between PLWH and controls, and enabled Omicron neutralization, 13% of SARS-CoV-2 naïve PLWH still no nAbs after 3 vaccine doses. This suggests additional doses/targeted vaccines could be merited, especially given 28% of SARS-CoV-2 naïve PLWH failed to neutralize Omicron after the third vaccine dose. Previous studies among similar cohorts of PLWH with undetectable HIV viral loads have produced mixed results, as previously reviewed (Mullender et al., 2022). SARS-CoV-2 viral vector vaccines have shown similar magnitude and durability of antibody responses to HIV-negative controls (Frater et al., 2021; Ogbe et al., 2022b) but reduced levels of seroconversion and neutralization have been reported after two doses in PLWH in a more recent study (Woldemeskel et al., 2022). Furthermore, viral vector vaccines, lower CD4 T cell count/viraemia and old age have been linked to lower serological responses and breakthrough infection (Sun et al., 2022). In terms of mRNA vaccines, both non-significant (Heftdal et al., 2022; Levy et al., 2021) and significant decreases in humoral responses have been reported in PLWH (Bessen et al., 2022; Brumme et al., 2022; Hensley et al., 2022; Jedicke et al., 2022). These differences may be due to the size of cohorts examined and the range of immune reconstitution in these PLWH. In contrast to previous work (Hassold et al., 2022; Nault et al., 2021; Noe et al., 2021; Touizer et al., 2021), we have found no association between the CD4 T cell count and serological outcome, which could be due to insufficient power in this study to detect differences. Moreover, few studies have addressed T cell activity after a third SARS-CoV-2 vaccine dose in this population, but in agreement with our findings a strong boosting effect of a third vaccine dose has been reported regardless of the CD4 T cell count (Vergori et al., 2022). Thus, the lower level of nAbs observed here in PLWH could be in part due to potential differences in boosting of memory responses to enable breadth against Omicron after three vaccine doses.

Serological data correlated significantly with frequency of spike-specific MBCs. The B cell phenotyping confirmed the characteristic and persistent defects seen in global MBCs in the setting of HIV (reviewed in (Moir and Fauci, 2017)). Specifically, we have observed lower frequencies of resting MBCs and higher frequencies of atypical and activated MBCs. This dysregulated MBC phenotype was also associated with a delay in developing nAbs after the first dose regardless of HIV status. Further evaluation of antigen-specific MBCs in a group of individuals after the third vaccine dose led to the interesting observation that spike-specific MBCs present in PLWH had a similar memory B cell phenotype as HIV-negative controls, albeit fewer resting MBCs. However, higher levels of global atypical MBCs, also observed in PLWH with lower neutralization at the third vaccine dose, suggest that the excess atypical MBCs may be effectively exhausted, as has been described (Moir et al., 2008). Therefore, SARS-CoV-2 serum antibody responses may be lower not because spike-specific responses are enriched within atypical MBCs and therefore unable to progress to an antibody secreting phenotype (as has been postulated for HIV/HBV (Burton et al., 2018; Meffre et al., 2016)), but rather because of global MBC disturbance. Thus, we propose that this reduced nAb to vaccination in PLWH may not be due to an alteration in the phenotype of antigen-specific cells but rather limited numbers of MBCs available to participate in the antigen-specific response via the canonical pathway.

In contrast to serological responses, SARS-CoV-2 vaccination elicited comparable T cell responses between PLWH and HIV-negative controls at all sampling points, and these responses were largely preserved against circulating VOCs, including Omicron, following three vaccine doses. Similarly, to the scenario seen in antibody responses, prior SARS-CoV-2 infection also resulted in higher T cell responses to vaccination (Lozano-Ojalvo et al., 2021; Prendecki et al., 2021; Reynolds et al., 2021). Interestingly, detectable T cell responses were noted in a proportion of SARS-CoV-2 naïve individuals at baseline (Alrubayyi et al., 2021; Ogbe et al., 2022a), which could represent pre-existing cross-reactive T cell cells due to past infection with other coronaviruses (Casado et al., 2022). An association between CD4 T cell counts and the magnitude of T cell responses was observed in SARS-CoV-2 naïve PLWH following vaccination, highlighting the relevance of immune cell reconstitution in producing effective immunity to vaccination, especially in people who lack memory responses elicited by natural infection. In this cohort, PLWH were well-controlled on ART and had undetectable HIV viral loads. Both PLWH with, and without, prior SARS-CoV-2 exposure had similar median CD4 T cell counts (602 and 560 cells/µl, respectively) despite different serological outcomes. However, the full impact of HIV-related immunosuppression, in addition to other factors, including age, sex and presence of co-morbidities, in dampening effective and long-lived memory responses needs to be addressed in future larger prospective studies. It is possible that different vaccine schedules, i.e., homologous versus heterologous vaccination, could also account for the observed heterogeneity in cellular immune responses. A heterologous viral vectored/mRNA vaccination has been described to lead to increased reactogenicity, combining the advantages from both vaccine classes (Banki et al., 2022). Due to limited numbers, it has not been possible to address the impact of different vaccine platforms in our cohort. Whether a heterologous approach induces more effective, resilient, and durable responses in PLWH merits further investigation to gain better insight into the design of the most effective/optimized vaccination schedules.

Overall humoral responses correlated with the magnitude of T cell responses and our findings corroborate the importance of T_FH_ cells supporting effective B cell responses after vaccination. Notably, in a small subgroup of patients (serological non-or low-level responders), there were detectable T cell responses characterised by a CXCR3+CD127+ CD8 T phenotype. This phenotype was not clearly related to HIV parameters or presence of co-morbidities. These T cell populations have been linked with increased survival in people infected with SARS-CoV-2 and are consistent with observations in patient groups who lack B cell responses (Gao et al., 2022). Upregulation of CXCR3 in vaccine-induced T cells with potential to home to lung mucosa in tuberculosis (Jeyanathan et al., 2017) suggests that these CD8 T cells described herein could play a role in the protection against severe respiratory diseases such as SARS-CoV-2. Future prospective studies in larger cohorts are needed to validate these findings and fully address how these vaccine-induced T cell responses could mediate protection, thereby guiding the design of novel immunization strategies.

Our study has several limitations. These include a cross-sectional analysis, which precludes the establishment of causal relationships. Our cohort is heterogeneous, with differences in sex, age and levels of immunosuppression that may contribute to the variability in the magnitude of responses. Moreover, the current analysis provides an overview of responses after up to three vaccine doses, and therefore further work is required to assess the durability and resilience of these responses against subvariants and additional vaccine doses.

Despite these caveats, our study provides new insights into the reasons why some PLWH fail to produce effective humoral responses, and an in-depth assessment of B cell responses. The observation of a more abundant CD8 T cell profile in some PLWH with absent or low-level antibody responses supports the notion that virus-specific CD8 T cells could compensate for defects in humoral immunity after SARS-CoV-2 vaccination in PLWH, as previously described for other immunocompromised groups (Gao et al., 2022). Overall, our data supports the benefit of a third SARS-CoV-2 dose in inducing nAbs against Omicron in PLWH, as it does in the general population. Future prospective studies are needed to fully evaluate humoral responses incorporating T cell metrics and potential early waning of responses to fully determine the correlates of protection against disease and the need for regular booster/altered vaccine schedules.

## Acknowledgements

We are grateful to all the clinic staff and participants at the Mortimer Market Centre and Ian Charleson Day Centre, in particular Rebecca Matthews, Aran Dhillon, Connor McAlpine, Paulina Prymas, Marzia Fiorino, Thomas Fernandez, Jonathan Edwards and Hemat Nargis. The authors would like to thank James E Voss of the Scripps Research Institute for the gift of Hela ACE2 expressing cells and Peter Cherepanov of the Francis Crick Institute for recombinant S1 antigen.

## Funding

This study was supported by UKRI MRC grant (MR/W020556/1) to LEM, DP and EM and an NIH award (R01AI55182) to (DP). LEM receives funding from the European Research Council (ERC) under the European Union’s 1053 Horizon 2020 research and innovation programme (Grant Agreement No. 757601) and is supported by a Career Development Award (MR/R008698/1). ET is supported by an MRC studentship (MR/N013867/1), AA by a Saudi Ministry of Education studentship (FG-350441) and TF by a Wellcome Trust Clinical PhD Fellowship (216358/Z/19/Z). NJM is supported by an MRC grant (MR/T032413/1), a NHSBT grant (WPA15-02), the Addenbrooke’s Charitable Trust (900239) and the NIHR Cambridge BRC.

## MATERIALS & METHODS

### Ethics statement

The protocols for the following study were approved by the local Research Ethics Committee (REC) Berkshire (REC 16/SC/0265) and South Central - Hampshire B (REC 19/SC/0423). The study complied with all relevant ethical regulations for work with human participants and conformed to the Helsinki declaration principles and Good Clinical Practice (GCP) guidelines. All subjects enrolled into the study provided written informed consent.

### Patient recruitment and sampling

There were 110 HIV+ participants who were virally suppressed and on ART and 64 HIV-negative healthy controls were recruited as part of either the Jenner II or the Vaccine in Clinical Infection (VCI) cohorts. PBMCs and plasma (or serum) were collected at the following timepoints: baseline, post-first dose (≥12 days following the first dose), post-second dose (≤70 days following the second dose), pre-third dose (≥70 days following the second dose), and post-third dose (>7 days following the third dose). Participants received a mix of available SARS-CoV-2 vaccination (Pfizer-BioNTech’s BNT162b2; Moderna’s mRNA-1273 or Astra-Zeneca’s AZD1222) according to Joint Committee on Vaccination and Immunization, UK, guidelines (Jcvi, 2022). Not every participant was sampled at all timepoints. At each visit, participants were asked to report any history of SARS-CoV-2 infection.

Between vaccinations, 4 previously SARS-CoV-2 naïve participants (2 HIV-, 2 HIV+) reported a SARS-CoV-2 infection, as such, any subsequent timepoints were moved into the ‘prior SARS-CoV-2 infection’ group for analysis. Similarly, 3 participants with prior SARS-CoV-2 reported a further infection (2 HIV-, 1 HIV+). All participants were recruited at the Mortimer Market Centre for Sexual Health and HIV Research and the Ian Charleson Day Centre at the Royal Free Hospital (London, UK) following written informed consent as part of a study approved by the local ethics board committee. Additional information about demographic and sampling can be found in Supplementary Table 1.

### PBMC isolation

Whole blood was collected in heparin-coated tubes. PBMCs were isolated from whole blood via density-gradient sedimentation. Whole blood was first spun via centrifugation for 5 min at 800g. Plasma was then collected, aliquoted and stored at −80°C for further use. Remaining blood was diluted with RPMI (Gibco), layered over an appropriate volume of Ficoll (Cytiva) and then spun via centrifugation for 20min at 800g without brake. The PBMC layer was collected and washed with RPMI to be spun via centrifugation for 10 min at 400g. PBMCs were stained with trypan blue and counted using Automated Cell Counter (BioRad, Hercules, California, USA). PBMCs were then cryopreserved in a cryovial in cell recovery freezing medium containing 10% dimethyl sulfoxide (DMSO) (Sigma) and 90% heat-inactivated fetal bovine serum (FBS) and stored at −80 °C in a Mr. Frosty freezing container overnight before being transferred into liquid nitrogen for further storage. If present, serum separator tubes were spun at 400g for 5 min to collect serum and then stored at −80°C for further use.

### Semi-quantitative S1 ELISA

This assay was set up previously by our lab (Alrubayyi et al., 2021; O’Nions et al., 2020). Briefly, in a 96-half-well NUNC Maxisorp™ plate (Nalgene, NUNC International, Hereford, UK), three columns were coated overnight at 4°C with 25 µl of goat anti-human F(ab)ʹ2 (1:1000) in PBS, the other nine columns were coated with 25µl of SARS-CoV-2 WT S1 protein (a kind gift from Peter Cherepanov (Ng et al., 2020), The Francis Crick Institute) at 3 µg/ml in PBS. The next day, plates were washed with PBS-T (0.05% Tween in PBS) and blocked for 1 hour (h) at room temperature (RT) with assay buffer (5% milk powder PBS-T). Assay buffer was then removed and 25 µl of patient plasma at dilutions from 1:50−1:10000 in assay buffer added to the S1-coated wells in duplicate. Serial dilutions of known concentrations of IgG were added to the F(ab)ʹ2 IgG-coated wells in triplicate to generate an internal standard curve. After 2 h of incubation at RT, plates were washed with PBS-T and 25 µl alkaline phosphatase (AP)-conjugated goat anti-human IgG (Jackson ImmunoResearch) at a 1:1000 dilution was added to each well and incubated for 1 h at RT. Plates were then washed with PBS-T, and 25 µl of AP substrate (Sigma Aldrich) added. Optical density (OD) was measured using a Multiskan™ FC (Thermo Fisher-Scientific UK) plate reader at 405 nm and S1-specific IgG titers were interpolated from the IgG standard curve using 4PL regression curve-fitting on GraphPad Prism 9.

### Total IgG ELISA

To measure total IgG levels in plasma, a 96-half-well NUNC Maxisorp™ plate (Nalgene, NUNC International, Hereford, UK) was entirely coated overnight at 4°C with 25 µl of goat anti-human F(ab)ʹ2 (1:1000). As above, plates were washed in PBS-T and blocked for 1 h at RT in assay buffer. 25µl of serial dilutions of patient plasma (1:100 to 1:10000000) were added in duplicates to the plate alongside known concentrations of IgG in triplicates. As above, after 2 h of incubation at RT, plates were washed with PBS-T and 25 µl AP-conjugated goat anti-human IgG was added and then incubated for 1 h at RT. Plates were washed with PBS-T, and 25 µl of AP substrate added. ODs were measured using a Multiskan™ FC (Thermo Fischer Scientific-UK)plate reader at 405 nm and total IgG titers interpolated from the IgG standard curve using 4PL regression curve-fitting on GraphPad Prism 9.

### IgG purification

As the PLWH participants in this study were on ART which can interfere with the lentivirus-based pseudotype neutralization assay IgG was purified from plasma using a 96-well protein G spin plate (Pierce™). Plasma was incubated in wells containing protein G at RT for 30 min. The captured IgG was then eluted with 0.1M Glycine (pH=2-3) twice into 2M Tris (pH=7.5-9) buffer. To remove Tris/Glycine buffer from the purified IgG, the eluate was concentrated (Thermo Scientific™ Pierce Protein Concentrator PES, 50K MWCO, 0.5 mL) and washed thrice at 10000rpm for 10 min before quantification by measuring absorbance of 280nm on a NanoDrop™ (ThermoFischer). The entire volume of purified IgG was then filtered sterile using a 0.22µm PDVF hydrophilic membrane FiltrEX™ filter plate (Corning) and stored at 4°C for further use.

### Pseudovirus production

In a T75 flask, 3×10^6^ HEK-293T cells were seeded in 10ml of complete DMEM Dulbecco’s Modified Eagle’s Medium (Gibco) supplemented with 10% FBS and 50μg/ml penicillin-streptomycin. The next day, the following transfection mix was prepared: 1ml of Opti-MEM™ (Gibco); 90µl of PEI-max (1mg/ml); 10µg of p8.91 HIV-1 gag/pol packaging plasmid (Zufferey et al., 1997); pCSLW HIV-1 luciferase reporter vector plasmid (Wright et al., 2008) and 5µg of either SARS-CoV-2 spike plasmid of interest, specifically WT (Wuhan-hu-1) or Omicron (BA.1/B.1.1.529.1) (Seow et al., 2020) as indicated in the results section. The transfection mix was left to incubate for 20 min before being added to the cells and left to incubate at 37°C 5% CO_2_ for 72h before being collected and filtered through a 0.45µm filter (Millipore) and either used directly in an assay or stored at −80°C.

### Pseudovirus neutralization

Neutralization assays were performed in 96-well plates by adding either duplicate serial dilutions of neat plasma in complete Dulbecco’s Modified Eagle medium (Thermo Fisher Scientific-UK (DMEM) starting at 1:20 dilution for HIV-negative samples or the appropriate amount of purified IgG for HIV+ samples to give a starting dilution equivalent to 200 or 400µg/ml of IgG as based on total IgG. These dilutions were incubated with the appropriate amount of filtered pseudotyped virus for 1h at 37°C 5% CO_2_ before adding 10000/ml HeLa-ACE2 cells (kind gift from James Voss, The Scripps Research Institute, USA) in 100µl per well. After a 72h incubation at 37°C 5% CO_2_, the supernatant was removed, and cells lysed. Bright-Glo™ luciferase substrate (Promega) was added, and relative light unit (RLU) values were read on a Glomax® (Promega) or BioTek Synergy™ H1 (Agilent) plate reader. RLU readouts were used to calculate the reciprocal inhibitory dilution at which 50% of the virus activity is neutralized by plasma (ID_50_) for each sample on GraphPad Prism 9.

### Live neutralization

The SARS-CoV-2 virus used in this study was the wildtype (lineage B) isolate SARS-CoV-2/human/Liverpool/REMRQ0001/2020, a kind gift from Ian Goodfellow (University of Cambridge), isolated by Lance Turtle (University of Liverpool) and David Matthews and Andrew Davidson (University of Bristol) (Daly et al., 2020; Patterson et al., 2020)(Plasma was heat-inactivated at 56°C for 30 mins before use, and neutralizing antibody titres at 50% inhibition (NT_50_) measured as previously described (Bergamaschi et al., 2021; Gerber et al., 2022; van der Klaauw et al., 2022). In brief, luminescent HEK293T-ACE2-30F-PLP2 reporter cells (clone B7) expressing SARS-CoV-2 Papain-like protease-activatable circularly permuted firefly luciferase (FFluc) were seeded in flat-bottomed 96-well plates. The next day, SARS-CoV-2 viral stock (MOI=0.01) was pre-incubated with a 3-fold dilution series of each sample for 2 h at 37°C, then added to the cells. 16 h post-infection, cells were lysed in Bright-Glo™ Luciferase Buffer (Promega) diluted 1:1 with PBS and 1% NP-40, and FFluc activity measured by luminometry. Experiments were conducted in duplicate. To obtain NT_50_, titration curves were plotted as FFluc vs log (serum dilution), then analysed by non-linear regression using the Sigmoidal, 4PL, X is log(concentration) function in GraphPad Prism. NT_50_ were quantitated when (1) at least 50% inhibition was observed at the lowest serum dilution tested (1:10, or 1:20 for pre-diluted samples), and (2) a sigmoidal curve with a good fit was generated. Samples with no detectable neutralizing activity were assigned an arbitrary NT_50_ equivalent to the lower limit of quantification.

### Production of biotinylated protein

To produce biotinylated spike and receptor binding domain (RBD) protein, HEK-293F cells were seeded at 1×10^6^ cells/ml in Freestyle™ 293 Expression Medium (Gibco). The next day, a transfection mix was prepared (for 200ml of cells) of 72µg of spike-Avi-His tag or RBD-Avi-His tag plasmid and 18µg of BirA plasmid (Graham et al., 2021; Seow et al., 2020) into 11ml of Opti-MEM™, alongside 2ml of PEI-Max® and 3ml of 10mM biotin, and left to incubate at 37°C 5% CO_2_ in a shaking incubator for 7 days before harvesting for purification. The supernatant was purified using an imidazole (Sigma-Aldrich) buffer at a final concentration of 20mM during binding to the His GraviTrap™ (Cytiva) column and 500mM imidazole for elution. The eluted protein was then concentrated with a 100KD Amicon® Ultra concentrator (Merck) and washed with PBS before quantification using a NanoDrop™. Biotinylated protein was then further purified through size exclusion chromatography using an AKTA™ pure system with a Superdex® 200 Increase 10/300 GL column (Sigma-Aldrich) to select for fractions containing trimeric spike or RBD protein.

### B cell phenotypic flow cytometric analysis

As previously described (Jeffery-Smith et al., 2022), 1µg of biotinylated spike with either streptavidin-conjugated allophycocyanin (APC) (ProZyme) and phycoerythrin (PE) (ProZyme) and 0.5µg of biotinylated RBD with BV421 (BioLegend) were incubated for 30 minutes in the dark to generate fluorochrome-linked biotinylated tetramers. Previously cryopreserved aliquots of 5×10^6^ or 10×10^6^ cell aliquots of PBMCs were quickly thawed in PBS, then stained with a panel of phenotyping antibodies and biotinylated tetramers (see Supplementary Table 2) or phenotyping antibodies only for FMO controls. PBMCs were then washed with PBS and fixed in Cytofix/Cytoperm™ (BD) buffer. Compensation controls were prepared according to manufacturer’s instructions using Anti-Mouse Ig, κ CompBeads™ (BD). Samples were acquired on an LSRFortessa™ (BD) flow cytometer. Data was analysed (see Supplementary Figure 2 for gating strategy) on FlowJo v10 (FlowJo, BD). For further analysis of the phenotype of spike-specific MBCs, analysis was limited to samples for which at least 50 cells were acquired in the CD19+ CD20+ CD38lo/-IgD-MBCs (excluding CD27-CD27+ cells) spike-PE+ spike-APC+ gate as previously defined (Jeffery-Smith et al., 2022).

| Protein | Fluorochrome | 1X PBS |
| --- | --- | --- |
| 1µg purified biotinylated spike | 1µl streptavidin-conjugated APC | 7µl |
| 1µg purified biotinylated spike | 0.5µl streptavidin-conjugated APC | 7.5µl |
| 0.5µg purified biotinylated spike | 1µl streptavidin-conjugated BV421 | 8µl |

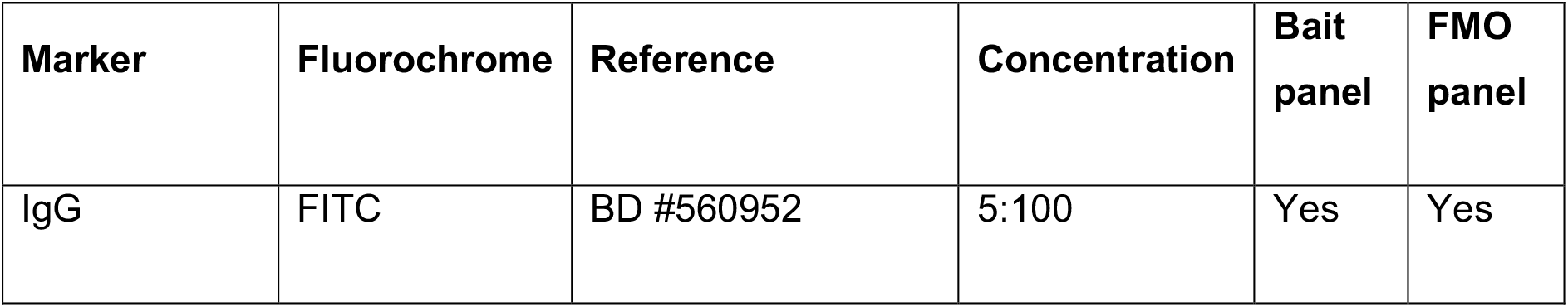
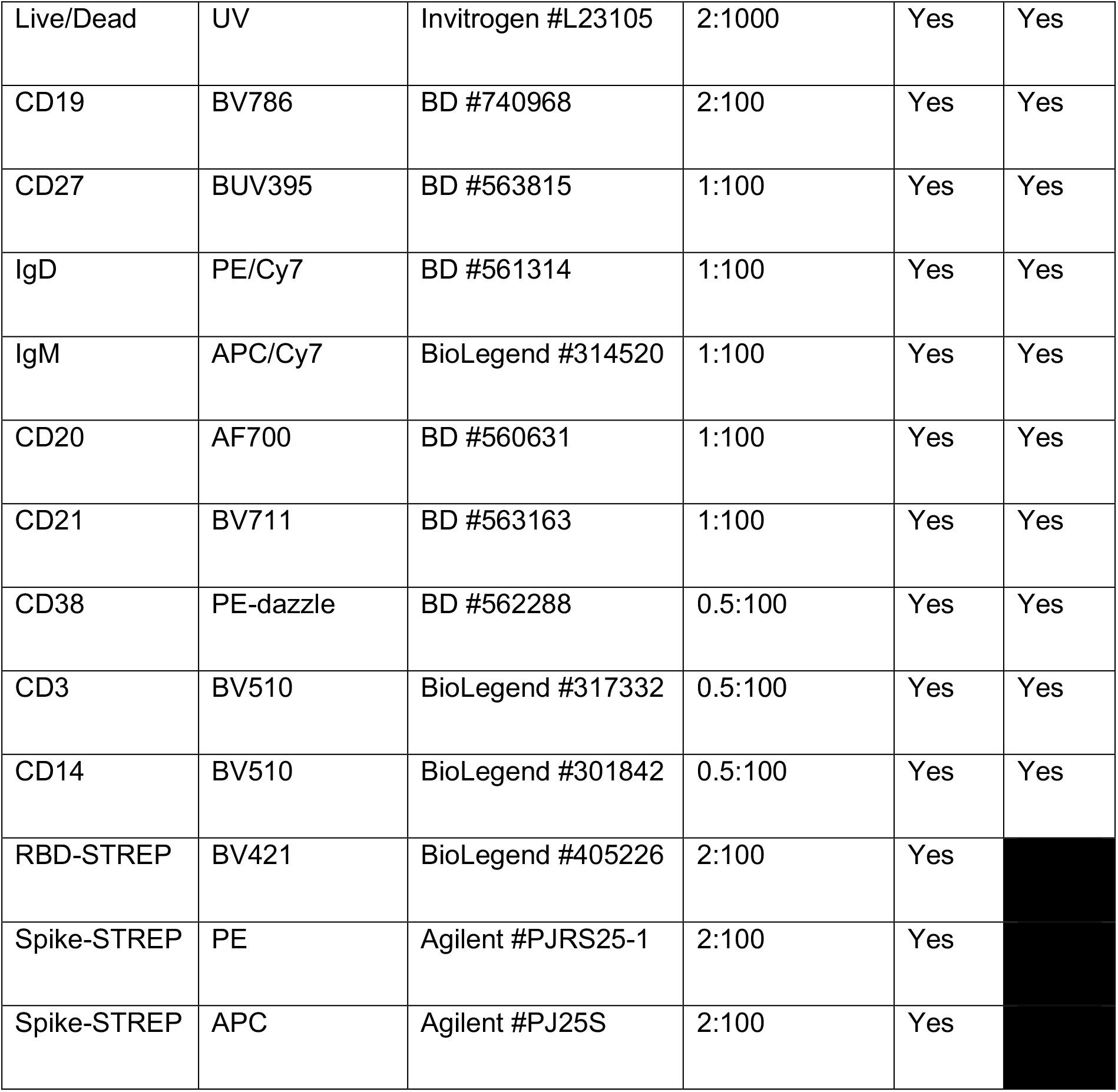

| Marker | Fluorochrome | Reference | Concentration | Bait panel | FMO panel |
| --- | --- | --- | --- | --- | --- |
| IgG | FITC | BD #560952 | 5:100 | Yes | Yes |
| Live/Dead | UV | Invitrogen #L23105 | 2:1000 | Yes | Yes |
| CD19 | BV786 | BD #740968 | 2:100 | Yes | Yes |
| CD27 | BUV395 | BD #563815 | 1:100 | Yes | Yes |
| IgD | PE/Cy7 | BD #561314 | 1:100 | Yes | Yes |
| IgM | APC/Cy7 | BioLegend #314520 | 1:100 | Yes | Yes |
| CD20 | AF700 | BD #560631 | 1:100 | Yes | Yes |
| CD21 | BV711 | BD #563163 | 1:100 | Yes | Yes |
| CD38 | PE-dazzle | BD #562288 | 0.5:100 | Yes | Yes |
| CD3 | BV510 | BioLegend #317332 | 0.5:100 | Yes | Yes |
| CD14 | BV510 | BioLegend #301842 | 0.5:100 | Yes | Yes |
| RBD-STREP | BV421 | BioLegend #405226 | 2:100 | Yes |  |
| Spike-STREP | PE | Agilent #PJRS25-1 | 2:100 | Yes |  |
| Spike-STREP | APC | Agilent #PJ25S | 2:100 | Yes |  |

### T cell phenotypic flow cytometric analysis

The flow cytometric analysis has been described in detail previously (Alrubayyi et al., 2021). Briefly, purified cryopreserved PBMCs samples were thawed and rested for 2 hours at 37°C in complete RPMI medium (RPMI supplemented with penicillin-streptomycin, L-Glutamine, HEPES, non-essential amino acids, 2-Mercaptoethanol, and 10% FBS). After 2-hour incubation, cells were washed and plated in a 96-round bottom plate at 0.5-1×10^6^ per well and stained for chemokine markers (CXCR3, CCR7 and CXCR5) for 30 minutes at 37 °C. Cells were then washed and stained with surface markers at 4 °C for 20 min with different combinations of antibodies in the presence of fixable live/dead stain (Invitrogen). After 20 min of incubation, cells were washed with PBS, and fixed with 4% paraformaldehyde for 15 min at RT. Samples were acquired on a LSRFortessa™ X-20 using FACSDiva™ version 8.0 (BD Biosciences) and subsequent data analysis was performed using FlowJo v10 (Treestar). The gating strategies used for flow cytometry experiments are provided in **Figure 6** and **Supplementary Figure 6**.

| Antibodies | Supplier | Identifier | Clone | Dilution |
| --- | --- | --- | --- | --- |
| APC/Cy7 anti-human CD197 (CCR7) | BioLegend | Cat # 353212 | Clone # G043H7 | 1 in 50ul |
| Brilliant Violet 650™ anti-human CD127 (IL-7Rα) Antibody | BioLegend | Cat # 351325 | Clone # A019D5 | 1 in 100ul |
| Brilliant Violet 650™ anti-human CD3 Antibody | BioLegend | Cat # 317324 | Clone # OKT3 | 1 in 100ul |
| Brilliant Violet 711™ anti-human CD27 Antibody | BioLegend | Cat # 302833 | Clone # O323 | 1 in 100ul |
| Brilliant Violet 785™ anti-human CD38 Antibody | BioLegend | Cat # 303530 | Clone # HIT2 | 1 in 50ul |
| Alexa Fluor® 700 anti-human CD45RA Antibody | BioLegend | Cat # 304120 | Clone # HI100 | 1 in 50ul |
| Brilliant Violet 421™ anti-human CD279 (PD-1) Antibody | BioLegend | Cat # 329920 | Clone # EH12.2H7 | 1 in 100ul |
| PE/Dazzle™ 594 anti-human CD4 Antibody | BioLegend | Cat # 300548 | Clone # RPA-T4 | 1 in 100ul |
| Brilliant Violet 711™ anti-human CD8a Antibody | BioLegend | Cat # 301044 | Clone # RPA-T8 | 1 in 100ul |
| Brilliant Violet 510™ anti-human CD14 Antibody | BioLegend | Cat # 301842 | Clone # M5E2 | 1 in 200ul |
| Brilliant Violet 510™ anti-human CD19 Antibody | BioLegend | Cat # 302242 | Clone # HIB19 | 1 in 200ul |
| BB515 Rat Anti-Human CXCR5 (CD185) | BD Biosciences | Cat # 564624 | Clone # RF8B2 | 1 in 50ul |
| BV605 Mouse Anti-Human CD56 | BD Biosciences | Cat # 562780 | Clone # NCAM16.2 | 1 in 50ul |
| PE-Cy7 Mouse Anti-Human CD25 | BD Biosciences | Cat # 335824 | Clone # 2A3 | 1 in 50ul |
| PE-Cy™5 Mouse Anti-Human CD183 | BD Biosciences | Cat # 551128 | Clone # 1C6/CXCR3 | 1 in 25ul |
| PerCP-eFluor 710 Anti-Human CD3 | eBioscience | Cat # 46-0037-42 | Clone # OKT3 | 1 in 100ul |
| APC Anti-Human CD19 | BioLegend | Cat # 302212 | Clone # HIB19 | 1 in 100ul |

### High-dimensional data analysis of flow cytometry data

Visualization of high-dimensional single-cell data (viSNE) (van der Maatens, 2014) and FlowSOM (Van Gassen et al., 2015) analyses were performed using the Cytobank platform (https://www.cytobank.org). Concatenated files were used to evaluate overall CD4 and CD8 T cell landscape in different groups. Cells were manually gated for lymphocytes, singlets, CD14-CD19-live cells, CD3+ and CD4+ or CD8+ and then subjected to viSNE analysis. The viSNE clustering analysis was performed on 8 parameters (CCR7, CD45RA, CD127, PD-1, CD38, CXCR5, CXCR3, CD25). Equal event sampling was selected across all samples. FlowSOM was then performed using the same markers outlined previously for viSNE and with the following parameters: number of clusters 100, number of metaclusters 10; the size of clusters 15 pixels (Cytobank default).

### Ex vivo IFN-γ ELISpot assay

The IFN-γ ELISpot assays were performed as described previously (Alrubayyi et al., 2021). Briefly, 96-well ELISpot plates (S5EJ044I10; Merck Millipore, Darmstadt, Germany) pre-wetted with 30 µl of 70% ethanol for 2 min before washing with 200 µl of sterile PBS. Anti-IFN-γ coating antibody (10 µg/ml in PBS; clone 1-D1K; Mabtech, Nacka Strand, Sweden) was then added and the plates incubated overnight at 4 °C. Prior to addition of cells, ELISpot plates were washed with PBS and blocked with R10 (RPMI supplemented with penicillin-streptomycin, L-glutamine, and 10% FBS) for a minimum of 2 h at 37 °C. Cells were then added at 2 × 10^5^ cells/well, in duplicate, and stimulated with overlapping peptide pools at 2 μg/ml for 16−18 h at 37 °C. Unstimulated cells were used as a negative control while PHA (10 µg/ml, Sigma-Aldrich) stimulated cells were used as a positive control. Plates were then washed with 0.05% Tween/PBS (Sigma Aldrich) and incubated for 2 h with an IFN-γ detection antibody (1 μg/ml; clone mAb-7B6-1; Mabtech) followed by 1 h incubation with AP-conjugated streptavidin (1:1000 in PBS, Mabtech). Plates were then washed and visualized using the VECTASTAIN® Elite® ABC-HRP kit according to the manufacturer’s instructions (Mabtech). Antigen-specific T cell responses were quantified by subtracting the number of spots in unstimulated cells from the peptide stimulated cells. An additional threshold of >5 SFU/10^6^ PBMCs was used. Participants who lacked T cell responses to the positive stimuli (PHA) or where antigen-specific responses found to be lower than two standard deviations of negative controls were excluded from the results.

### Overlapping peptide pools

For the detection of antigen-specific T cell responses, purified cryopreserved PBMCs were stimulated with the following peptide pools: (1) Wild-type SARS-CoV-2 spike; SARS-CoV-2 spike PepTivator® protein pools (Miltenyi Biotec, Gladbach, GER) were used to test T cell responses against full spike proteome. (2) VOC spike-specific peptide pools; the WuhanHu-1 and variant pools containing peptides from the Wuhan Hu-1, Alpha (B.1.1.7), Beta (B.1.351), Delta (B.1.617.2) and Omicron (BA.1/B.1.1.529.1) sequences (9, 19, 32 and 83 peptides, respectively) were used to define T cell responses to mutated Spike sequences in SARS-CoV-2 variants. Alpha and Beta peptide pools were synthesised by GL Biochem Shanghai Ltd, China and previously used in (Reynolds et al., 2021). The corresponding controls to Alpha and Beta pools with Wuhan Hu-1amino-acid sequences were compared in parallel within the same donor. Delta and Omicron pools were obtained from Miltenyi Biotec. (3) Non-SARS-CoV-2 antigens: Peptide pools of the pp65 protein of human cytomegalovirus (CMV) (Miltenyi Biotec, Gladbach, GER), or HIV-1 gag peptide pools (NIH AIDS Reagent Repository) were used as positive/negative controls.

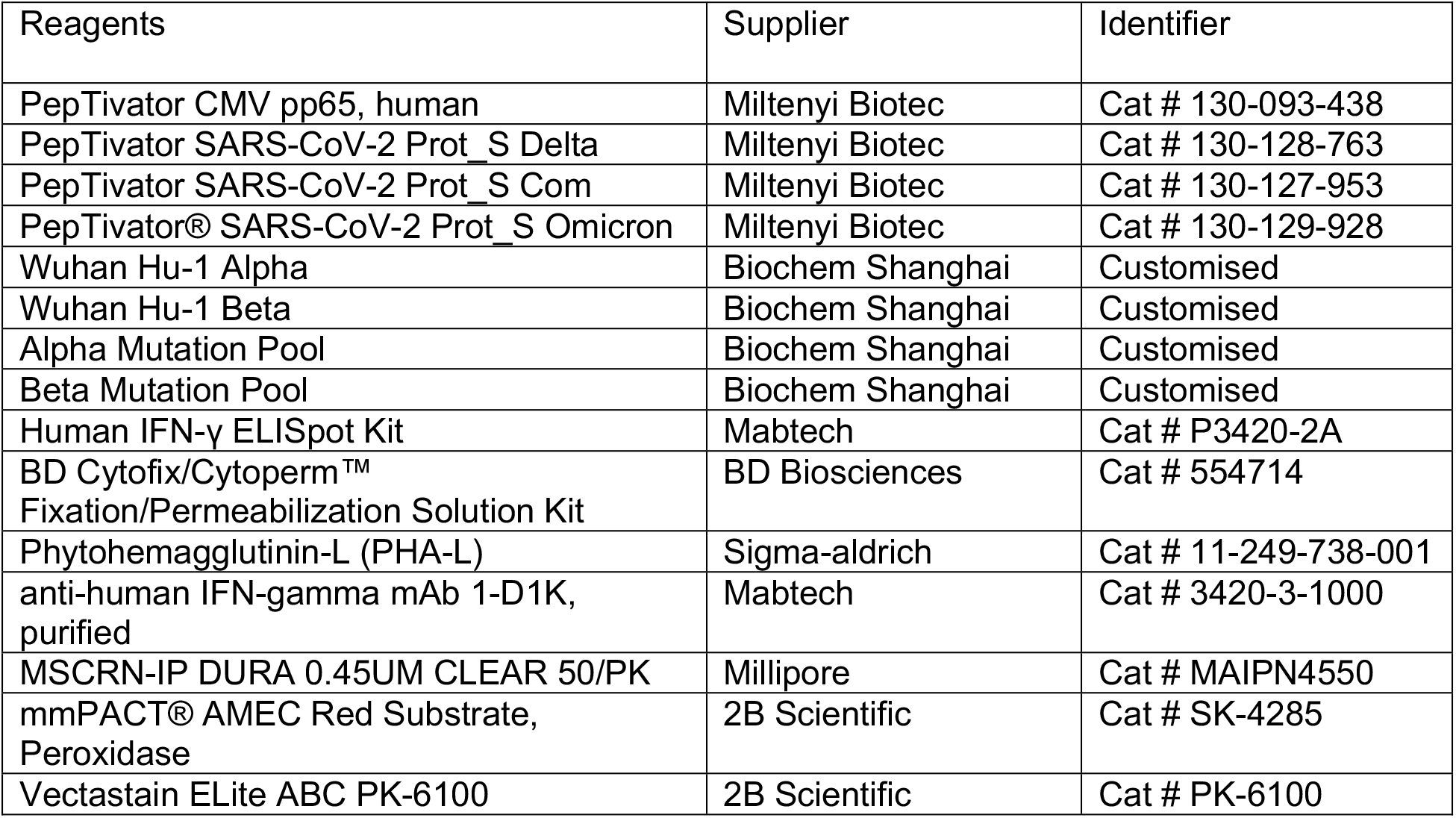

### Statistical analysis

All statistical analysis were carried out in GraphPad Prism 9.0 (GraphPad). All tests were two-tailed. Mann-Whitney U-test (MWU) was used to compare unpaired, non-parametric data whilst Wilcoxon matched-pairs sign rank test (WMP) was used to compare paired, non-parametric data. Non-parametric Spearman test was used for correlation analysis between two sets of data. Error bars represent mean + standard deviation. Statistical significance in the figures is shown as p-value >0.0332 (*); >0.0021 (**); >0.0002 (***) and >0.0001 (****).

## Notes

### Competing Interest Statement

The authors have declared no competing interest.

## REFERENCES

Adam, L., Rosenbaum, P., Quentric, P., Parizot, C., Bonduelle, O., Guillou, N., Corneau, A., Dorgham, K., Miyara, M., Luyt, C.E., et al. (2021). CD8+PD-L1+CXCR3+ polyfunctional T cell abundances are associated with survival in critical SARS-CoV-2-infected patients. JCI Insight 6.

Alrubayyi, A., Gea-Mallorqui, E., Touizer, E., Hameiri-Bowen, D., Kopycinski, J., Charlton, B., Fisher-Pearson, N., Muir, L., Rosa, A., Roustan, C., et al. (2021). Characterization of humoral and SARS-CoV-2 specific T cell responses in people living with HIV. Nat Commun 12, 5839.

Banki, Z., Mateus, J., Rossler, A., Schafer, H., Bante, D., Riepler, L., Grifoni, A., Sette, A., Simon, V., Falkensammer, B., et al. (2022). Heterologous ChAdOx1/BNT162b2 vaccination induces stronger immune response than homologous ChAdOx1 vaccination: The pragmatic, multi-center, three-arm, partially randomized HEVACC trial. EBioMedicine 80, 104073.

Baskaran, V., Lawrence, H., Lansbury, L.E., Webb, K., Safavi, S., Zainuddin, N.I., Huq, T., Eggleston, C., Ellis, J., Thakker, C., et al. (2021). Co-infection in critically ill patients with COVID-19: an observational cohort study from England. J Med Microbiol 70.

Bergamaschi, L., Mescia, F., Turner, L., Hanson, A.L., Kotagiri, P., Dunmore, B.J., Ruffieux, H., De Sa, A., Huhn, O., Morgan, M.D., et al. (2021). Longitudinal analysis reveals that delayed bystander CD8+ T cell activation and early immune pathology distinguish severe COVID-19 from mild disease. Immunity 54, 1257–1275 e1258.

Bertagnolio, S., Thwin, S.S., Silva, R., Nagarajan, S., Jassat, W., Fowler, R., Haniffa, R., Reveiz, L., Ford, N., Doherty, M., et al. (2022). Clinical features of, and risk factors for, severe or fatal COVID-19 among people living with HIV admitted to hospital: analysis of data from the WHO Global Clinical Platform of COVID-19. Lancet HIV 9, e486–e495.

Bessen, C., Plaza-Sirvent, C., Bhat, J., Marheinecke, C., Urlaub, D., Bonowitz, P., Busse, S., Schumann, S., Vidal Blanco, E., Skaletz-Rorowski, A., et al. (2022). Impact of SARS-CoV-2 vaccination on systemic immune responses in people living with HIV. medRxiv, 2022.2004.2008.22273605-22272022.22273604.22273608.22273605.

Braun, J., Loyal, L., Frentsch, M., Wendisch, D., Georg, P., Kurth, F., Hippenstiel, S., Dingeldey, M., Kruse, B., Fauchere, F., et al. (2020). SARS-CoV-2-reactive T cells in healthy donors and patients with COVID-19. Nature 587, 270–274.

Brouwer, P.J.M., Caniels, T.G., van der Straten, K., Snitselaar, J.L., Aldon, Y., Bangaru, S., Torres, J.L., Okba, N.M.A., Claireaux, M., Kerster, G., et al. (2020). Potent neutralizing antibodies from COVID-19 patients define multiple targets of vulnerability. Science 369, 643–650.

Brumme, Z.L., Mwimanzi, F., Lapointe, H.R., Cheung, P.K., Sang, Y., Duncan, M.C., Yaseen, F., Agafitei, O., Ennis, S., Ng, K., et al. (2022). Humoral immune responses to COVID-19 vaccination in people living with HIV receiving suppressive antiretroviral therapy. NPJ Vaccines 7.

Burton, A.R., Pallett, L.J., McCoy, L.E., Suveizdyte, K., Amin, O.E., Swadling, L., Alberts, E., Davidson, B.R., Kennedy, P.T., Gill, U.S., et al. (2018). Circulating and intrahepatic antiviral B cells are defective in hepatitis B. J Clin Invest 128, 4588–4603.

Casado, J.L., Vizcarra, P., Haemmerle, J., Velasco, H., Martin-Hondarza, A., Rodriguez-Dominguez, M.J., Velasco, T., Martin, S., Romero-Hernandez, B., Fernandez-Escribano, M., et al. (2022). Pre-existing T cell immunity determines the frequency and magnitude of cellular immune response to two doses of mRNA vaccine against SARS-CoV-2. Vaccine X 11, 100165.

Cohen, K.W., Linderman, S.L., Moodie, Z., Czartoski, J., Lai, L., Mantus, G., Norwood, C., Nyhoff, L.E., Edara, V.V., Floyd, K., et al. (2021). Longitudinal analysis shows durable and broad immune memory after SARS-CoV-2 infection with persisting antibody responses and memory B and T cells. Cell Rep Med 2, 100354.

Cooper, T.J., Woodward, B.L., Alom, S., and Harky, A. (2020). Coronavirus disease 2019 (COVID-19) outcomes in HIV/AIDS patients: a systematic review. HIV Med 21, 567–577.

Cruciani, M., Mengoli, C., Serpelloni, G., Lanza, A., Gomma, M., Nardi, S., Rimondo, C., Bricolo, F., Consolaro, S., Trevisan, M.T., et al. (2009). Serologic response to hepatitis B vaccine with high dose and increasing number of injections in HIV infected adult patients. Vaccine 27, 17–22.

Daly, J.L., Simonetti, B., Klein, K., Chen, K.E., Williamson, M.K., Anton-Plagaro, C., Shoemark, D.K., Simon-Gracia, L., Bauer, M., Hollandi, R., et al. (2020). Neuropilin-1 is a host factor for SARS-CoV-2 infection. Science 370, 861–865.

Dan, J.M., Mateus, J., Kato, Y., Hastie, K.M., Yu, E.D., Faliti, C.E., Grifoni, A., Ramirez, S.I., Haupt, S., Frazier, A., et al. (2020). Immunological memory to SARS-CoV-2 assessed for up to eight months after infection. bioRxiv.

Dandachi, D., Geiger, G., Montgomery, M.W., Karmen-Tuohy, S., Golzy, M., Antar, A.A.R., Llibre, J.M., Camazine, M., Diaz-De Santiago, A., Carlucci, P.M., et al. (2021). Characteristics, Comorbidities, and Outcomes in a Multicenter Registry of Patients With Human Immunodeficiency Virus and Coronavirus Disease 2019. Clin Infect Dis 73, e1964–e1972.

Fenwick, C., Joo, V., Jacquier, P., Noto, A., Banga, R., Perreau, M., and Pantaleo, G. (2019). T-cell exhaustion in HIV infection. Immunol Rev 292, 149–163.

Frater, J., Ewer, K.J., Ogbe, A., Pace, M., Adele, S., Adland, E., Alagaratnam, J., Aley, P.K., Ali, M., Ansari, M.A., et al. (2021). Safety and immunogenicity of the ChAdOx1 nCoV-19 (AZD1222) vaccine against SARS-CoV-2 in HIV infection: a single-arm substudy of a phase 2/3 clinical trial. The Lancet HIV 0.

Fuster, F., Vargas, J.I., Jensen, D., Sarmiento, V., Acuna, P., Peirano, F., Fuster, F., Arab, J.P., Martinez, F., and Core, H.I.V.S.G. (2016). CD4/CD8 ratio as a predictor of the response to HBV vaccination in HIV-positive patients: A prospective cohort study. Vaccine 34, 1889–1895.

Gao, Y., Cai, C., Wullimann, D., Niessl, J., Rivera-Ballesteros, O., Chen, P., Lange, J., Cuapio, A., Blennow, O., Hansson, L., et al. (2022). Immunodeficiency syndromes differentially impact the functional profile of SARS-CoV-2-specific T cells elicited by mRNA vaccination. Immunity.

George, V.K., Pallikkuth, S., Parmigiani, A., Alcaide, M., Fischl, M., Arheart, K.L., and Pahwa, S. (2015). HIV infection Worsens Age-Associated Defects in Antibody Responses to Influenza Vaccine. The Journal of Infectious Diseases 211, 1959–1959.

Gerber, P.P., Duncan, L.M., Greenwood, E.J., Marelli, S., Naamati, A., Teixeira-Silva, A., Crozier, T.W., Gabaev, I., Zhan, J.R., Mulroney, T.E., et al. (2022). A protease-activatable luminescent biosensor and reporter cell line for authentic SARS-CoV-2 infection. PLoS Pathog 18, e1010265.

Gilbert, P.B., Montefiori, D.C., McDermott, A.B., Fong, Y., Benkeser, D., Deng, W., Zhou, H., Houchens, C.R., Martins, K., Jayashankar, L., et al. (2022). Immune correlates analysis of the mRNA-1273 COVID-19 vaccine efficacy clinical trial. Science 375, 43–50.

Goel, R.R., Painter, M.M., Apostolidis, S.A., Mathew, D., Meng, W., Rosenfeld, A.M., Lundgreen, K.A., Reynaldi, A., Khoury, D.S., Pattekar, A., et al. (2021). mRNA vaccines induce durable immune memory to SARS-CoV-2 and variants of concern. Science 374.

Graham, C., Seow, J., Huettner, I., Khan, H., Kouphou, N., Acors, S., Winstone, H., Pickering, S., Galao, R.P., Dupont, L., et al. (2021). Neutralization potency of monoclonal antibodies recognizing dominant and subdominant epitopes on SARS-CoV-2 Spike is impacted by the B.1.1.7 variant. Immunity 54, 1–14.

Grifoni, A., Weiskopf, D., Ramirez, S.I., Mateus, J., Dan, J.M., Moderbacher, C.R., Rawlings, S.A., Sutherland, A., Premkumar, L., Jadi, R.S., et al. (2020). Targets of T Cell Responses to SARS-CoV-2 Coronavirus in Humans with COVID-19 Disease and Unexposed Individuals. Cell 181, 1489–1501 e1415.

Hassold, N., Brichler, S., Ouedraogo, E., Leclerc, D., Carroue, S., Gater, Y., Alloui, C., Carbonnelle, E., Bouchaud, O., Mechai, F., et al. (2022). Impaired antibody response to COVID-19 vaccination in advanced HIV infection. AIDS 36, F1–F5.

Heftdal, L.D., Knudsen, A.D., Hamm, S.R., Hansen, C.B., Møller, D.L., Pries-Heje, M., Fogh, K., Hasselbalch, R.B., Jarlhelt, I., Pérez-Alós, L., et al. (2022). Humoral response to two doses of BNT162b2 vaccination in people with HIV. Journal of Internal Medicine 291, 513–513.

Hensley, K.S., Jongkees, M.J., Geers, D., GeurtsvanKessel, C.H., Ash Dalm, V., Papageorgiou, G., Steggink, H., Gorska, A., den Hollander, J.G., Bierman, W.F.W., et al. (2022). Immunogenicity and reactogenicity of SARS-CoV-2 vaccines in people living with HIV: a nationwide prospective cohort study in the Netherlands. medRxiv, 2022.2003.2031.22273221-22272022.22273203.22273231.22273221.

Hoffmann, C., Casado, J.L., Harter, G., Vizcarra, P., Moreno, A., Cattaneo, D., Meraviglia, P., Spinner, C.D., Schabaz, F., Grunwald, S., et al. (2021). Immune deficiency is a risk factor for severe COVID-19 in people living with HIV. HIV Med 22, 372–378.

Jcvi (2022). COVID-19 Greenbook chapter 14a.

Jedicke, N., Stankov, M.V., Cossmann, A., Dopfer-Jablonka, A., Knuth, C., Ahrenstorf, G., Ramos, G.M., and Behrens, G.M.N. (2022). Humoral immune response following prime and boost BNT162b2 vaccination in people living with HIV on antiretroviral therapy. HIV Medicine 23, 558–563.

Jeffery-Smith, A., Burton, A.R., Lens, S., Rees-Spear, C., Davies, J., Patel, M., Gopal, R., Muir, L., Aiano, F., Doores, K.J., et al. (2022). SARS-CoV-2–specific memory B cells can persist in the elderly who have lost detectable neutralizing antibodies. Journal of Clinical Investigation 132.

Jeyanathan, M., Afkhami, S., Khera, A., Mandur, T., Damjanovic, D., Yao, Y., Lai, R., Haddadi, S., Dvorkin-Gheva, A., Jordana, M., et al. (2017). CXCR3 Signaling Is Required for Restricted Homing of Parenteral Tuberculosis Vaccine-Induced T Cells to Both the Lung Parenchyma and Airway. J Immunol 199, 2555–2569.

Kernéis, S., Launay, O., Turbelin, C., Batteux, F., Hanslik, T., and Boëlle, P.Y. (2014). Long-term immune responses to vaccination in HIV-infected patients: A systematic review and meta-analysis. Clinical Infectious Diseases 58, 1130–1139.

Le Bert, N., Tan, A.T., Kunasegaran, K., Tham, C.Y.L., Hafezi, M., Chia, A., Chng, M.H.Y., Lin, M., Tan, N., Linster, M., et al. (2020). SARS-CoV-2-specific T cell immunity in cases of COVID-19 and SARS, and uninfected controls. Nature 584, 457–462.

Levy, I., Wieder-Finesod, A., Litchevsky, V., Biber, A., Indenbaum, V., Olmer, L., Huppert, A., Mor, O., Goldstein, M., Levin, E.G., et al. (2021). Immunogenicity and safety of the BNT162b2 mRNA COVID-19 vaccine in people living with HIV-1. Clinical Microbiology and Infection 27, 1851–1851.

Lozano-Ojalvo, D., Camara, C., Lopez-Granados, E., Nozal, P., Del Pino-Molina, L., Bravo-Gallego, L.Y., Paz-Artal, E., Pion, M., Correa-Rocha, R., Ortiz, A., et al. (2021). Differential effects of the second SARS-CoV-2 mRNA vaccine dose on T cell immunity in naive and COVID-19 recovered individuals. Cell Rep 36, 109570.

Mateus, J., Grifoni, A., Tarke, A., Sidney, J., Ramirez, S.I., Dan, J.M., Burger, Z.C., Rawlings, S.A., Smith, D.M., Phillips, E., et al. (2020). Selective and cross-reactive SARS-CoV-2 T cell epitopes in unexposed humans. Science 370, 89–94.

McCallum, M., Czudnochowski, N., Rosen, L.E., Zepeda, S.K., Bowen, J.E., Walls, A.C., Hauser, K., Joshi, A., Stewart, C., Dillen, J.R., et al. (2022). Structural basis of SARS-CoV-2 Omicron immune evasion and receptor engagement. Science 375, 864–868.

Meffre, E., Louie, A., Bannock, J., Kim, L.J.Y., Ho, J., Frear, C.C., Kardava, L., Wang, W., Buckner, C.M., Wang, Y., et al. (2016). Maturational characteristics of HIV-specific antibodies in viremic individuals. JCI Insight 1,: e84610-:e84610.

Moir, S., and Fauci, A.S. (2013). Insights Into B Cells And Hiv-Specific B-Cell Responses In Hiv-Infected Individuals. Immunological Reviews 254, 207–224.

Moir, S., and Fauci, A.S. (2017). B-cell responses to HIV infection (Blackwell Publishing Ltd), pp. 33–48.

Moir, S., Ho, J., Malaspina, A., Wang, W., DiPoto, A.C., O’Shea, M.A., Roby, G., Kottilil, S., Arthos, J., Proschan, M.A., et al. (2008). Evidence for HIV-associated B cell exhaustion in a dysfunctional memory B cell compartment in HIV-infected viremic individuals. Journal of Experimental Medicine 205, 1797–1805.

Mullender, C., da Costa, K.A.S., Alrubayyi, A., Pett, S.L., and Peppa, D. (2022). SARS-CoV-2 immunity and vaccine strategies in people with HIV. Oxford Open Immunology, iqac005.

Nasi, M., De Biasi, S., Gibellini, L., Bianchini, E., Pecorini, S., Bacca, V., Guaraldi, G., Mussini, C., Pinti, M., and Cossarizza, A. (2017). Ageing and inflammation in patients with HIV infection. Clinical and Experimental Immunology 187, 44–52.

Nault, L., Marchitto, L., Goyette, G., Tremblay-Sher, D., Fortin, C., Martel-Laferrière, V., Trottier, B., Richard, J., Durand, M., Kaufmann, D., et al. (2021). Covid-19 vaccine immunogenicity in people living with HIV-1. bioRxiv, 2021.2008.2013.456258-452021.456208.456213.456258.

Ng, K.W., Faulkner, N., Cornish, G.H., Rosa, A., Harvey, R., Hussain, S., Ulferts, R., Earl, C., Wrobel, A.G., Benton, D.J., et al. (2020). Preexisting and de novo humoral immunity to SARS-CoV-2 in humans. Science 370, 1339–1343.

Noe, S., Ochana, N., Wiese, C., Schabaz, F., Von Krosigk, A., Heldwein, S., Rasshofer, R., Wolf, E., and Jonsson-Oldenbuettel, C. (2021). Humoral response to SARS-CoV-2 vaccines in people living with HIV. Infection 1, 1–1.

Ogbe, A., Pace, M., Bittaye, M., Tipoe, T., Adele, S., Alagaratnam, J., Aley, P.K., Ansari, M.A., Bara, A., Broadhead, S., et al. (2022a). Durability of ChAdOx1 nCoV-19 vaccination in people living with HIV. JCI Insight 7.

Ogbe, A., Pace, M., Bittaye, M., Tipoe, T., Adele, S., Alagaratnam, J., Aley, P.K., Ansari, M.A., Bara, A., Broadhead, S., et al. (2022b). Durability of ChAdOx1 nCoV-19 vaccination in people living with HIV. JCI Insight 7.

Pallikkuth, S., De Armas, L.R., Pahwa, R., Rinaldi, S., George, V.K., Sanchez, C.M., Pan, L., Dickinson, G., Rodriguez, A., Fischl, M., et al. (2018). Impact of aging and HIV infection on serologic response to seasonal influenza vaccination. AIDS 32, 1085–1094.

Patterson, E.I., Prince, T., Anderson, E.R., Casas-Sanchez, A., Smith, S.L., Cansado-Utrilla, C., Solomon, T., Griffiths, M.J., Acosta-Serrano, A., Turtle, L., et al. (2020). Methods of Inactivation of SARS-CoV-2 for Downstream Biological Assays. J Infect Dis 222, 1462–1467.

Pensieroso, S., Galli, L., Nozza, S., Ruffin, N., Castagna, A., Tambussi, G., Hejdeman, B., Misciagna, D., Riva, A., Malnati, M., et al. (2013). B-cell subset alterations and correlated factors in HIV-1 infection. Aids 27, 1209–1217.

Portugal, S., Tipton, C.M., Sohn, H., Kone, Y., Wang, J., Li, S., Skinner, J., Virtaneva, K., Sturdevant, D.E., Porcella, S.F., et al. (2015). Malaria-associated atypical memory B cells exhibit markedly reduced B cell receptor signaling and effector function. Elife 4.

Prendecki, M., Clarke, C., Brown, J., Cox, A., Gleeson, S., Guckian, M., Randell, P., Pria, A.D., Lightstone, L., Xu, X.N., et al. (2021). Effect of previous SARS-CoV-2 infection on humoral and T-cell responses to single-dose BNT162b2 vaccine. Lancet 397, 1178–1181.

Rees-Spear, C., Muir, L., Griffith, S.A., Heaney, J., Aldon, Y., Snitselaar, J.L., Thomas, P., Graham, C., Seow, J., Lee, N., et al. (2021). The effect of spike mutations on SARS-CoV-2 neutralization. Cell Reports 34.

Reynolds, C.J., Pade, C., Gibbons, J.M., Butler, D.K., Otter, A.D., Menacho, K., Fontana, M., Smit, A., Sackville-West, J.E., Cutino-Moguel, T., et al. (2021). Prior SARS-CoV-2 infection rescues B and T cell responses to variants after first vaccine dose. Science.

Sekine, T., Perez-Potti, A., Rivera-Ballesteros, O., Stralin, K., Gorin, J.B., Olsson, A., Llewellyn-Lacey, S., Kamal, H., Bogdanovic, G., Muschiol, S., et al. (2020). Robust T Cell Immunity in Convalescent Individuals with Asymptomatic or Mild COVID-19. Cell 183, 158–168 e114.

Seow, J., Graham, C., Merrick, B., Acors, S., Pickering, S., Steel, K.J.A., Hemmings, O., O’Byrne, A., Kouphou, N., Galao, R.P., et al. (2020). Longitudinal observation and decline of neutralizing antibody responses in the three months following SARS-CoV-2 infection in humans. Nature Microbiology 5.

Spinelli, M.A., Peluso, M.J., Lynch, K.L., Yun, C., Glidden, D.V., Henrich, T.J., Deeks, S.G., and Gandhi, M. (2021). Differences in Post-mRNA Vaccination Severe Acute Respiratory Syndrome Coronavirus 2 (SARS-CoV-2) Immunoglobulin G (IgG) Concentrations and Surrogate Virus Neutralization Test Response by Human Immunodeficiency Virus (HIV) Status and Type of Vaccine: A Ma. Clinical Infectious Diseases.

Sun, J., Zheng, Q., Madhira, V., Olex, A.L., Anzalone, A.J., Vinson, A., Singh, J.A., French, E., Abraham, A.G., Mathew, J., et al. (2022). Association Between Immune Dysfunction and COVID-19 Breakthrough Infection After SARS-CoV-2 Vaccination in the US. JAMA Intern Med 182, 153–162.

Tamuzi, J.L., Muyaya, L.M., Mitra, A., and Nyasulu, P.S. (2022). Systematic review and meta-analysis of COVID-19 vaccines safety, tolerability, and efficacy among HIV-infected patients. medRxiv, 2022.2001.2011.22269049.

Terreri, S., Piano Mortari, E., Vinci, M.R., Russo, C., Alteri, C., Albano, C., Colavita, F., Gramigna, G., Agrati, C., Linardos, G., et al. (2022). Persistent B cell memory after SARS-CoV-2 vaccination is functional during breakthrough infections. Cell Host Microbe 30, 400–408 e404.

Touizer, E., Alrubayyi, A., Rees-Spear, C., Fisher-Pearson, N., Griffith, S.A., Muir, L., Pellegrino, P., Waters, L., Burns, F., Kinloch, S., et al. (2021). Failure to seroconvert after two doses of BNT162b2 SARS-CoV-2 vaccine in a patient with uncontrolled HIV. The Lancet HIV 8, e317–e318.

van der Klaauw, A.A., Horner, E.C., Pereyra-Gerber, P., Agrawal, U., Foster, W.S., Spencer, S., Vergese, B., Smith, M., Henning, E., Ramsay, I.D., et al. (2022). Accelerated waning of the humoral response to SARS-CoV-2 vaccines in obesity. medRxiv, 2022.2006.2009.22276196.

van der Maatens, L. (2014). Accelerating t-SNE using Tree-Based Algorithms. J Mach Learn Res 15, 3221–3245.

Van Gassen, S., Callebaut, B., Van Helden, M.J., Lambrecht, B.N., Demeester, P., Dhaene, T., and Saeys, Y. (2015). FlowSOM: Using self-organizing maps for visualization and interpretation of cytometry data. Cytometry A 87, 636–645.

Vergori, A., Cozzi Lepri, A., Cicalini, S., Matusali, G., Bordoni, V., Lanini, S., Meschi, S., Iannazzo, R., Mazzotta, V., Colavita, F., et al. (2022). Immunogenicity to COVID-19 mRNA vaccine third dose in people living with HIV. Nat Commun 13, 4922.

Western Cape Department of Health in collaboration with the National Institute for Communicable Diseases, S.A. (2021). Risk Factors for Coronavirus Disease 2019 (COVID-19) Death in a Population Cohort Study from the Western Cape Province, South Africa. Clin Infect Dis 73, e2005–e2015.

Woldemeskel, B.A., Karaba, A.H., Garliss, C.C., Beck, E.J., Wang, K.H., Laeyendecker, O., Cox, A.L., and Blankson, J.N. (2022). The BNT162b2 mRNA Vaccine Elicits Robust Humoral and Cellular Immune Responses in People Living With Human Immunodeficiency Virus (HIV). Clin Infect Dis 74, 1268–1270.

Wright, E., Temperton, N.J., Marston, D.A., McElhinney, L.M., Fooks, A.R., and Weiss, R.A. (2008). Investigating antibody neutralization of lyssaviruses using lentiviral pseudotypes: a cross-species comparison. The Journal of General Virology 89, 2204–2204.

Yang, X., Sun, J., Patel, R.C., Zhang, J., Guo, S., Zheng, Q., Olex, A.L., Olatosi, B., Weissman, S.B., Islam, J.Y., et al. (2021). Associations between HIV infection and clinical spectrum of COVID-19: a population level analysis based on US National COVID Cohort Collaborative (N3C) data. Lancet HIV 8, e690–e700.

Zufferey, R., Nagy, D., Mandel, R.J., Naldini, L., and Trono, D. (1997). Multiply attenuated lentiviral vector achieves efficient gene delivery in vivo. Nature Biotechnology 15, 871–875.

