## Supplementary information for "Attenuated humoral responses in HIV infection after SARS-CoV-2 vaccination are linked to global B cell defects and cellular immune profiles"

**Supplementary table 1 related to table 1: Cohort Demographics and Clinical Characteristics for PLWH SARS-CoV-2 naïve (nAb$^{-/low}$T$^{+}$ or nAb$^{+}$T$^{+}$).**

**Supplementary Figure 1: Lower post vaccination antibody responses in SARS-CoV-2 naïve PLWH.**

**Supplementary Figure 2: Gating strategy for spike-specific and global MBC.**

**Supplementary Figure 3: Improved neutralization against Omicron after the 3$^{rd}$ dose in PLWH accompanied by minimal alteration in spike-specific MBC phenotype.**

**Supplementary figure 4. Magnitude of T cell responses to Gag, CMV, and VOCs peptide pools and associations with age, days post vaccination and vaccine platform.**

**Supplementary figure. 5: Correlation between spike-specific T cell response and S1 IgG titers in HIV-positive and -negative individuals.**

**Supplementary figure 6. T cell immunophenotyping (or T cell differentiation) in HIV-negative and HIV-positive donors.**

**Supplementary table 1 related to table 1: Cohort Demographics and Clinical Characteristics for PLWH SARS-CoV-2 naïve (nAb$^{-/low}$T$^{+}$ or nAb$^{+}$T$^{+}$)**
Demographic and clinical characteristics of HIV-positive SARS-CoV-2 naïve individuals who lacked or had low nAbs ($ID_{50}$ <150) but had detectable virus-specific T cell responses following two doses of vaccination.

| | | HIV+SARS-CoV-2 naïve nAbs$^{low/-}$, T cell responses$^{+}$ | HIV+SARS-CoV-2 naïve nAbs$^{high}$, T cell responses$^{+}$ |
|---|---|---|---|
| Group size | n | 9 | 9 |
| COVID-19 vaccine | mRNA-based vaccine (BNT162b2/Pfizer or Moderna), n (%) | 3 (33.3%) | 5 (55.6%) |
| | ChAdOx1/ AstraZeneca, n (%) | 6 (66.7%) | 4 (44.4%) |
| | Day post 2$^{nd}$ dose, median (range) | 16 (12-32) | 19 (7-41) |
| Demographics | Age, median (range) | 52 (35-63) | 57 (29-62) |
| | Sex, n female:male | 1:8 | 0:9 |
| | Ethnicity, White: BAME | 7:2 | 7:2 |
| HIV parameters | HIV viral load | <50 | <50 |
| | CD4, median (range) | 680 (470-1360) | 650 (380-820) |
| | CD4:CD8, median (range) | 1 (0.4-3.05) | 1 (0.39-1.29) |
| Pre-existing conditions | None, n (%) | 7 (77.8%) | 6 (66.7%) |
| | Respiratory disease (asthma and COPD), n (%) | - | 1 (11.1%) |
| | Liver disease, n (%) | 1 (11.1%) | - |
| | Bone disease, n (%) | 1 (11.1%) | 3 (33.3%) |

**Supplementary Figure 1: Lower post vaccination antibody responses in SARS-CoV-2 naïve PLWH**

- **(A)** SARS-CoV-2 S1 IgG-specific responses (µg/ml) were measured by semiquantitative ELISA in PLWH in PLWH (blue) compared to HIV-negative controls (grey) stratified by vaccination timepoints for individuals without prior SARS-CoV-2 infection. The dotted line represents lower limit of the assay (0.6µg/ml), each data point is representative of n=2 biological repeats. N numbers match those in Figure 1A, Statistical test: MWU.
- **(B)** Shows the equivalent data for those with prior SARS-CoV-2 infection.
- **(C)** Correlation between WT $ID_{50}$ titres and S1 IgG-specific (µg/ml) titres stratified by PLWH (blue) and controls (grey) at all timepoints, statistical test: Spearman's correlation.
- **(D)** Correlation between WT $ID_{50}$ titres and live-SARS-CoV-2 (WT) $NT_{50}$ titres for PLWH (blue, n=32) and HIV-negative controls (grey, n=14). Dotted lines represent lower limits of both assays (1:20). Live-SARS-CoV-2 $NT_{50}$ represents a single biological repeat.
- **(E)** WT $ID_{50}$ titres in PLWH (blue) compared to HIV-negative controls (grey) stratified by vaccination timepoint for individuals without prior SARS-CoV-2 infection who received mRNA vaccines. Statistical test: MWU.
- **(F)** Shows the equivalent data for those with prior SARS-CoV-2 infection
- **(G)** WT $ID_{50}$ titres after at the 3$^{rd}$ dose for SARS-CoV-2 naïve PLWH (blue) stratified into either with or without comorbidities (see table for details) compared to HIV-negative controls (grey). Statistical test: MWU.
- **(H)** Longitudinal semi-quantitative ELISA titres for HIV-negative controls without prior SARS-CoV-2 infection who provided samples after the first and second vaccine dose and were categorised as exhibiting standard neutralizing response (coloured grey), or delayed neutralization if neutralization was only achieved after the second dose (colour-coded in magenta). N numbers for each category are indicated on the graph.
- **(I)** Shows the equivalent data for PLWH without prior SARS-CoV-2 infection
- **(J)** Shows the equivalent data for HIV-negative controls with prior SARS-CoV-2 infection
- **(K)** Shows the equivalent data for PLWH with prior SARS-CoV-2 infection
- **(L)** Correlation between WT $ID_{50}$ titres and S1 IgG-specific (µg/ml) titres stratified by standard (grey) or delayed neutralization (magenta) at all timepoints, statistical test: Spearman's correlation.
- **(M)** Correlation between WT $ID_{50}$ titres and CD4 count or
- **(N)** CD4:CD8 ratio stratified by standard (grey) or delayed neutralization (magenta) at all timepoints, statistical test: Spearman's correlation.

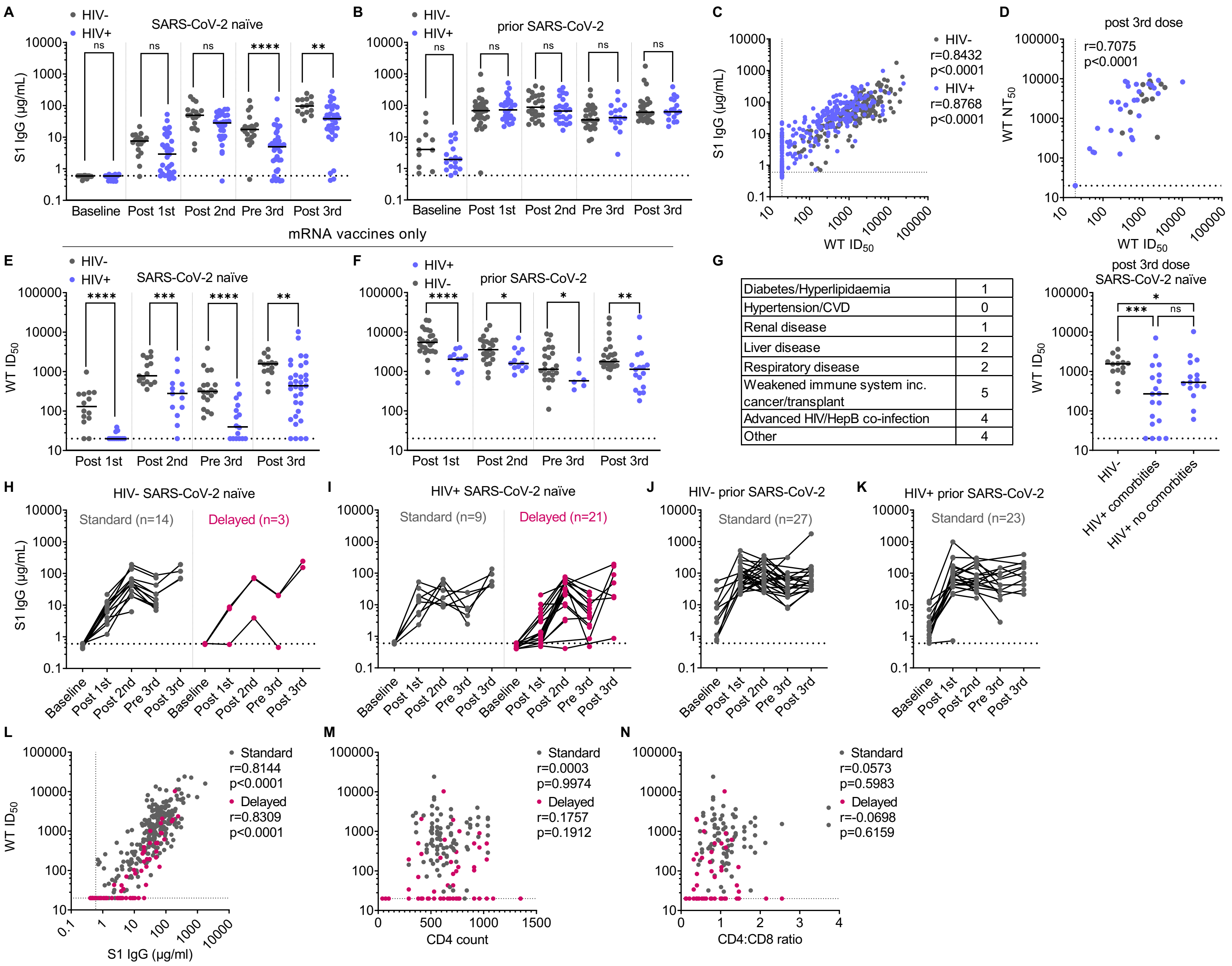


**Supplementary Figure 1 relating to Figure 1**

**Supplementary Figure 2: Gating strategy for spike-specific and global MBC**

Singlet live lymphocytes were first gated on, then using CD3/CD14 to gate out T cells and monocytes and CD19 to gate on B cells. Memory B cells were gated as CD20+ CD38$^{lo/-}$ cells and selecting for class-switched cells (i.e., IgD-). This allowed to further gate on specific MBC phenotype using CD21 and CD27. 'True' MBCs were defined by excluding the CD21+ CD27- switched naïve population to then gate on isotypes using IgG and IgM or spike-specific MBCs by using biotinylated baits for spike and RBD.

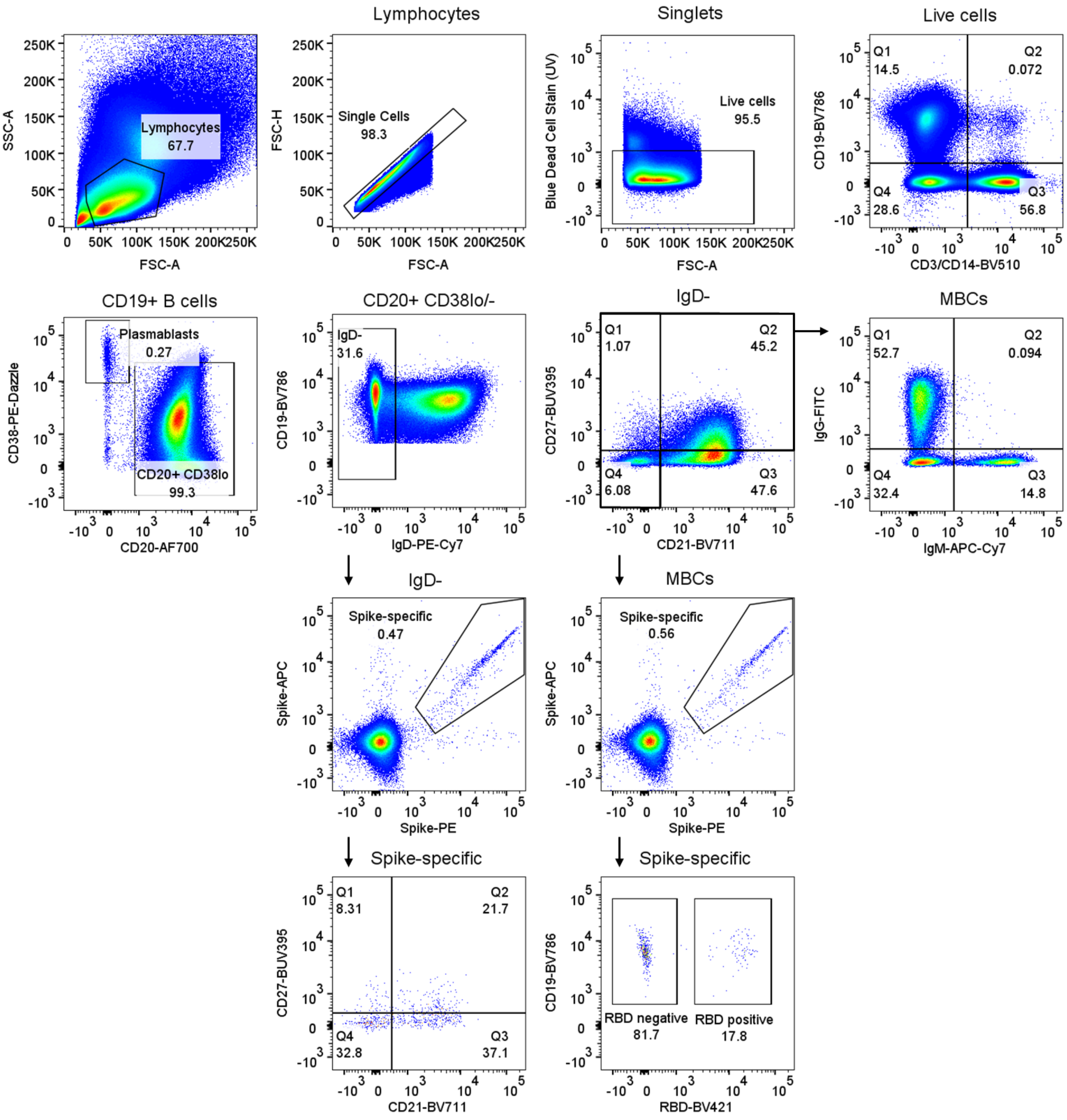


**Supplementary Figure 2 relating to Figure 2**

**Supplementary Figure 3: Improved neutralization against Omicron after the 3rd dose in PLWH accompanied by minimal alteration in spike-specific MBC phenotype**

**(A)** WT $ID_{50}$ titres in PLWH (blue) and HIV-negative donors (grey) included in the flow cytometry analysis after the 3rd dose stratified by SARS-CoV-2 infection at the 3rd dose. Statistical test: MWU.

**(B)** Shows the equivalent data as (A) for neutralisation against Omicron

**(C)** Omicron $ID_{50}$ titres after the 3rd dose for SARS-CoV-2 naïve PLWH (blue) stratified into either with or without comorbidities (see table in Figure 1G for details) compared to HIV-negative controls (grey).

**(D)** Frequency of spike-specific MBCs after the 3rd dose for SARS-CoV-2 naïve PLWH (blue) stratified into either with or without comorbidities (see table for details) compared to HIV-negative controls (grey).

**(E)** Correlation between donor age and WT $ID_{50}$ at post 1st, 2nd, 3rd and pre 3rd dose in SARS-CoV-2 naïve PLWH (blue) and HIV-negative controls (grey). Statistical test: Spearman's correlation.

**(F)** Correlation between donor age and days since last vaccine dose at post 1st, 2nd, 3rd and pre 3rd dose. Statistical test: in SARS-CoV-2 naïve PLWH (blue) and HIV-negative controls (grey). Spearman's correlation.

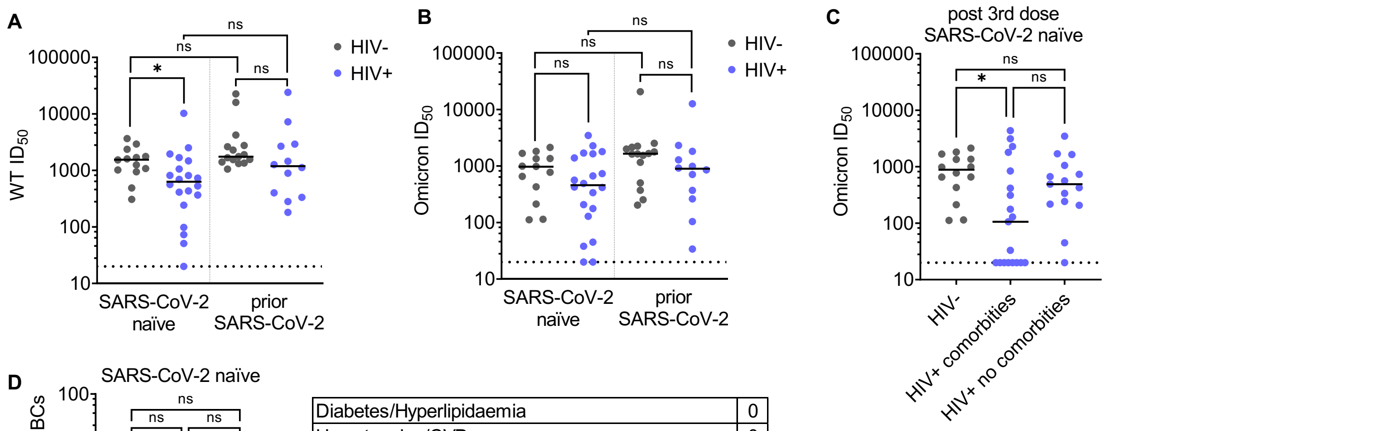


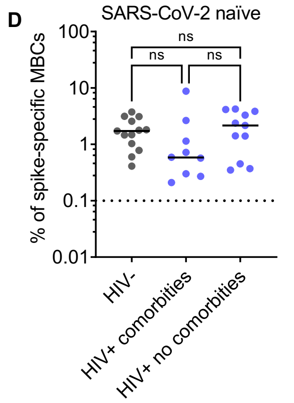


| | |
|---|---|
| Diabetes/Hyperlipidaemia | 0 |
| Hypertension/CVD | 0 |
| Renal disease | 0 |
| Liver disease | 2 |
| Respiratory disease | 1 |
| Weakened immune system inc. cancer/transplant | 3 |
| Advanced HIV/HepB co-infection | 1 |
| Other | 2 |

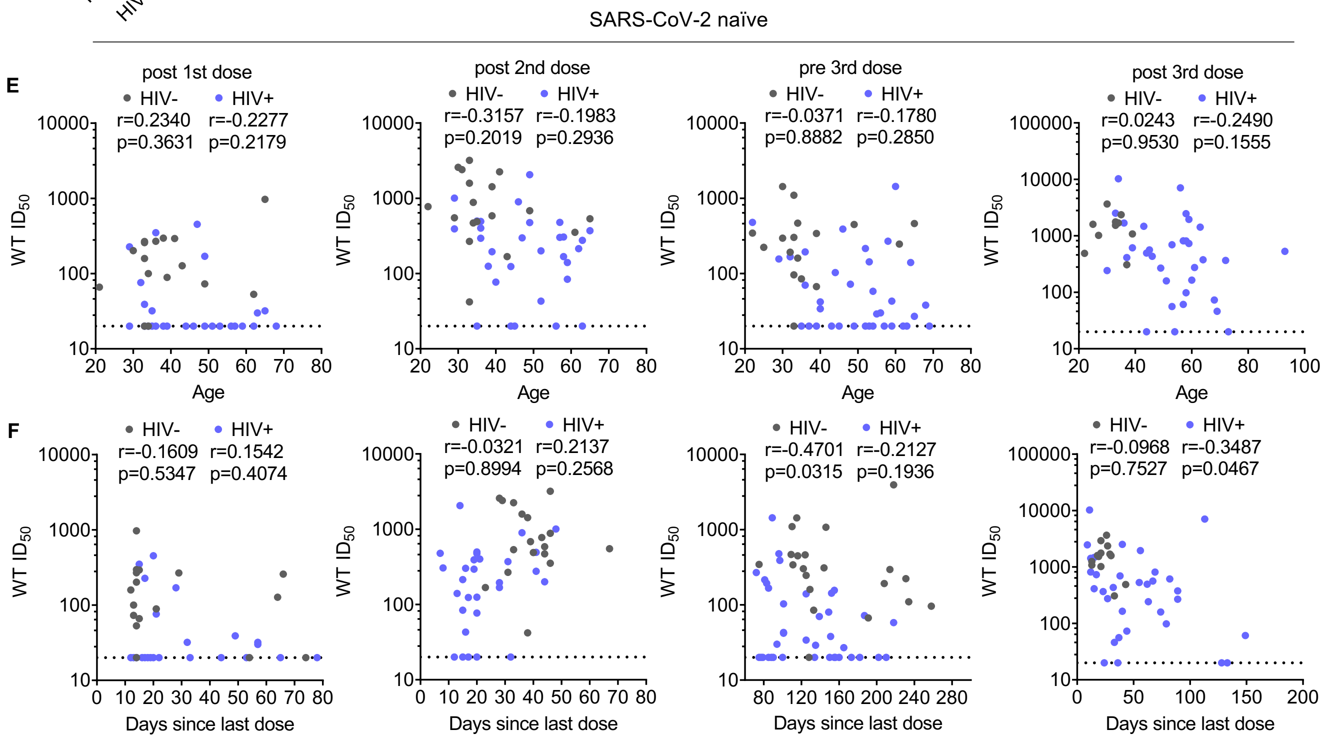


**Supplementary Figure 3 relating to Figure 3**

**Supplementary figure 4. Magnitude of T cell responses to Gag, CMV, and VOCs peptide pools and associations with age, days post vaccination and vaccine platform.**

**(A-C)** Paired analysis of the IFN-γ-ELISpot responses for SARS-CoV-2 (spike), CMV (pp65), HIV (Gag), and PHA in HIV-negative and HIV-positive donors after 1st dose (A), 2nd dose (B), and 3rd dose (C) of vaccine. Statistical test: Wilcoxon matched-pairs sign rank test (WMP).

**(D-E)** The magnitude of T cell responses to Wuhan, Alpha, Beta, and Delta after three doses of the vaccine in HIV-negative and HIV-positive donors with no prior SARS-CoV-2 infection (D) and with prior SARS-CoV-2 exposure (E). Statistical test: WMP.

**(F-H)** Correlation between CD4:CD8 ratio in HIV-infected individuals and the magnitude of spike-specific T cell responses after 1st dose (F), 2nd dose (G), and (H) 3rd dose. Statistical test: Spearman's rank correlation coefficient.

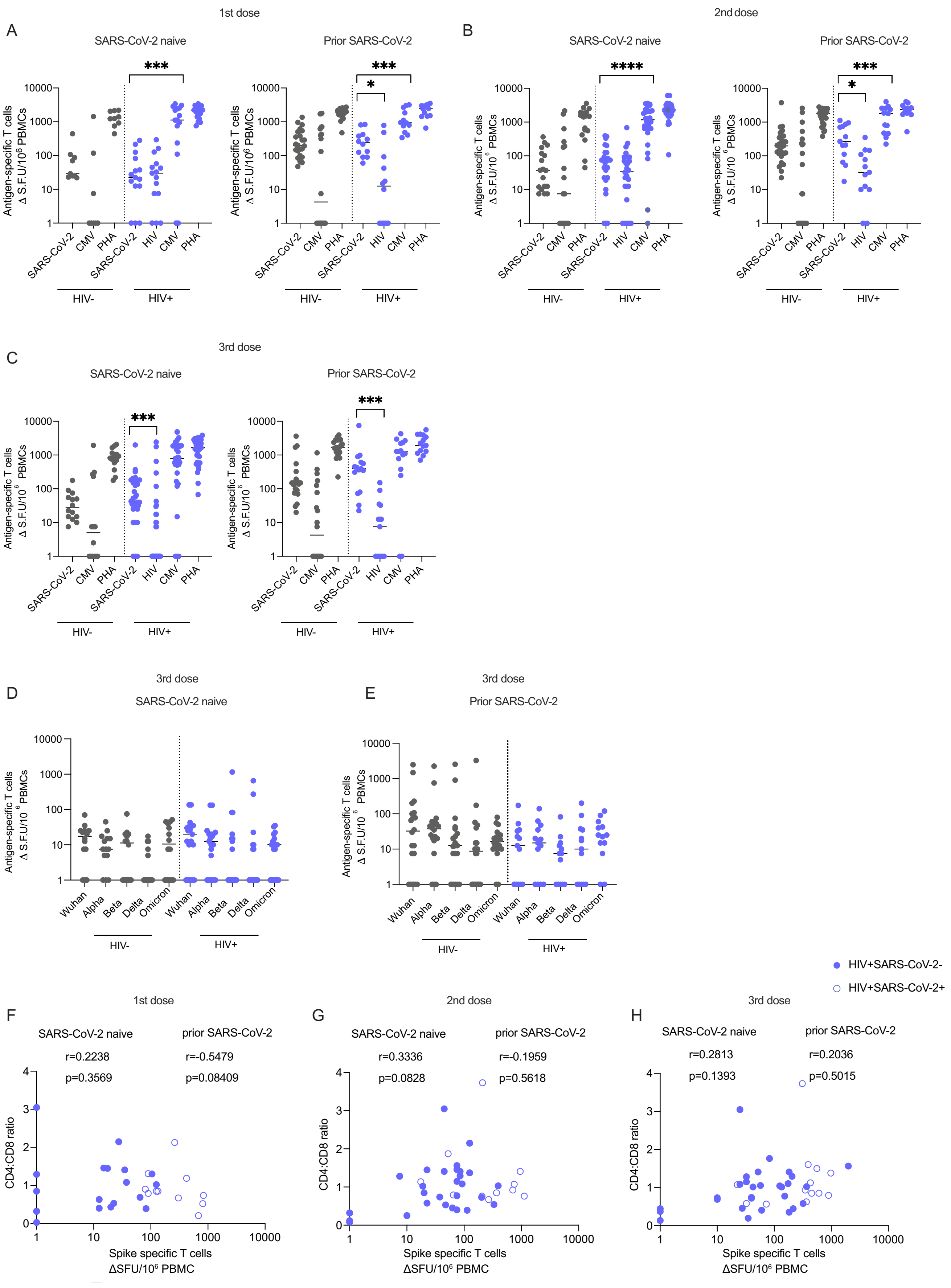


**Supplementary Figure 4 relating to Figure 4**

**Supplementary figure. 5: Correlation between spike-specific T cell response and S1 IgG titers in HIV-positive and -negative individuals.**

**(A-C)** Correlation of spike-specific T cell responses with S1 IgG titers after first dose (A), second dose (B) , and third dose (C) of vaccine in HIV-negative and HIV-positive donors, with or without prior SARS-CoV-2 infection (the limit of detection S1 IgG=0.6 µg/ml). Statistical test: Spearman's rank correlation coefficient.

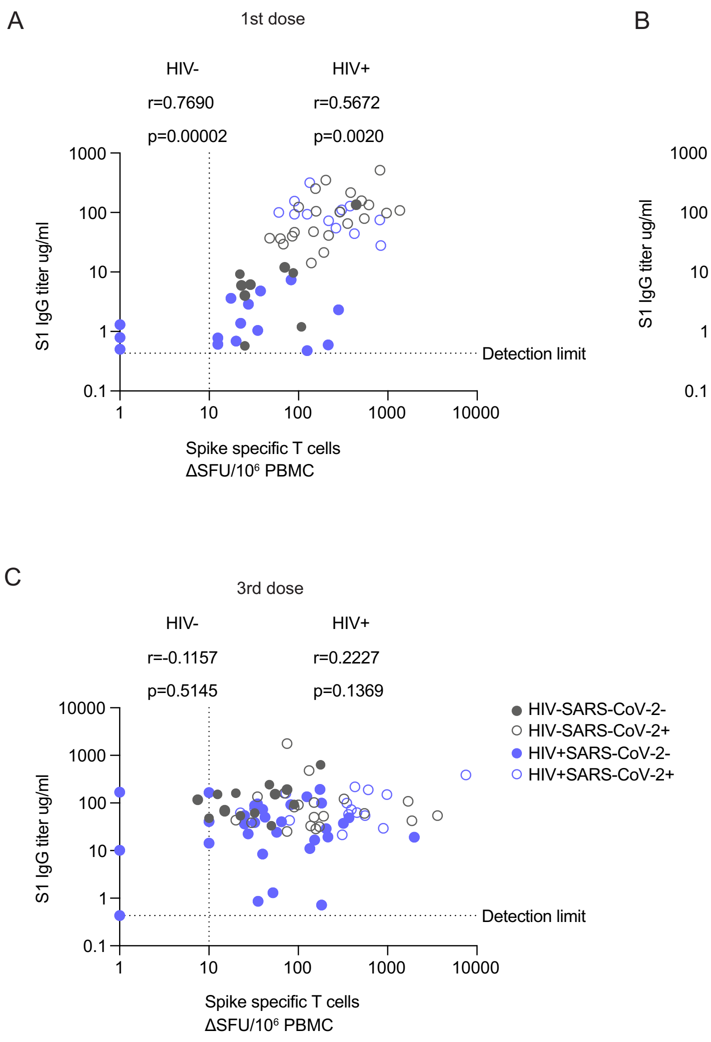


**Supplementary Figure 5 relating to Figure 5**

**Supplementary figure 6. T cell immunophenotyping (or T cell differentiation) in HIV-negative and HIV-positive donors**

- **(A)** Surface expression intensity heatmap of the markers indicated for each of the ten FlowSOM meta-clusters of CD4 T cells (showing in Figure. 6 A&B). (colour scale: row z-score expression for each individual marker).
- **(B)** Heatmap of the markers for CD8 T cell clusters (showing in Figure. 3 F&G).
- **(C)** Representative flow plots of the gating strategy for the identification of CM ($CD45RA^{-}/CCR7^{+}$ central memory), naïve ($CD45RA^{+}/CCR7^{+}$), TEMRA ($CD45RA^{+}/CCR7^{-}$ terminally differentiated effector memory) and EM ($CD45RA^{-}/CCR7^{-}$ effector memory) CD4 and CD8 T cells in HIV-negative and HIV-positive donors after two doses of the vaccine.
- **(D)** Summary analysis of the percentage of CD4 and CD8 T cell subsets in $nAb^{-/low}$ (opened blue circle) and $nAb^{high}$ (filled blue circle) individuals (n=9 in each group). Statistical test: Mann-Whitney U-test (MWU).
- **(E-F)** FlowSOM meta-clusters of CD4 (E) and CD8 (F) T cells from $nab^{-/low}$ and $nab^{high}$ HIV-positive SARS-CoV-2 naïve subjects after three doses of vaccine (n=5 $nAb^{-/low}$, n=13 $nAb^{high}$).

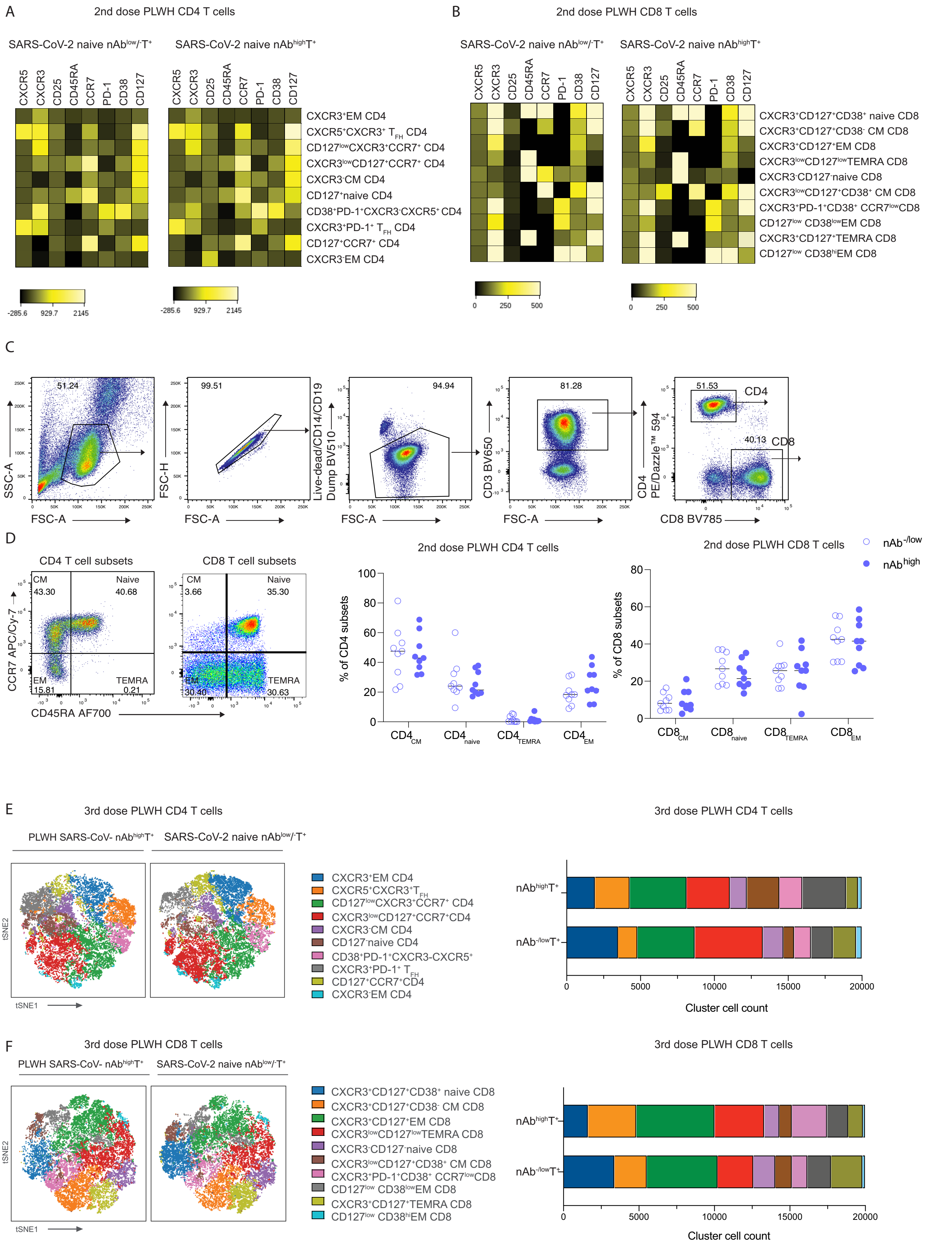


**Supplementary Figure 6 relating to Figure 6**
